# Serotonin, dopamine, and norepinephrine transporter assembly is selectively disrupted by a NET truncation isoform as revealed through near-million-atom simulations

**DOI:** 10.64898/2026.03.25.714186

**Authors:** Taner Karagöl, Alper Karagöl

## Abstract

**Background:** Monoamine transporters (MATs), including the dopamine, norepinephrine, and serotonin transporters (DAT, NET, SERT), are essential regulators of synaptic neurotransmission that rely on complex oligomeric bio-assemblies for proper function. The regulatory influence of naturally occurring, alternatively spliced truncated isoforms on bio-assembly dynamics remains profoundly underexplored.

**Methods:** To decode these interactions at the atomic level, we deployed a multiscale computational framework. We integrated genomics-guided multimer predictions with massive-scale, near-million-atom molecular dynamics (MD) simulations within explicit lipid bilayers. The thermodynamic stability of these heteromeric complexes was quantified using membrane-adapted MM/PBSA calculations, which were subsequently correlated with dynamics-aware evolutionary profiling to map co-evolutionary interaction hotspots.

**Results:** Our analyses reveal that the NET-derived truncated isoform A0A804HLI4 acts as a pan-family potential inhibitor. It forms stable, exergonic heterodimers with canonical NET and DAT, thermodynamically outcompeting native homodimerization. In full tetrameric simulations, the integration of a single isoform precipitates a macro-structural disruptions of the SERT complex. The variant anchors at non-native interfaces, locking the assembly into an asymmetric, non-native state. Residue-level thermodynamic decomposition and evolutionary mapping isolated conserved structural elements (most notably Gln236) that dictate this high-affinity cross-reactivity across the SLC6 family.

**Conclusions:** Truncated MAT isoforms execute a dynamic mechanism of inhibitory effects and may systematically downregulate synaptic reuptake capacity by sequestering functional monomers. These findings establish a thermodynamically grounded, high-resolution model of isoform-induced bio-assembly disruptions. Crucially, they expose these non-canonical, isoform-driven interfaces as conserved and highly druggable targets, offering a distinct pharmacological paradigm for precision interventions in neuropsychiatric and neurodegenerative pathologies.

## Introduction

Monoamine transporters (MATs), comprising the dopamine, norepinephrine, and serotonin transporters, are integral membrane proteins that govern neurochemical homeostasis by mediating the presynaptic reuptake of their respective neurotransmitters [1,2]. By facilitating the high-affinity reuptake of these neurotransmitters into the presynaptic terminal, MATs effectively terminate monoaminergic signaling and recycle substrates for subsequent release. The plasma membrane transporters: the norepinephrine transporter (NET), dopamine transporter (DAT), and serotonin transporter (SERT), are central to maintaining neurochemical homeostasis [3]. Consequently, dysregulation of these systems is a hallmark of diverse neuropsychiatric conditions including anxiety, attention-deficit hyperactivity disorder, and major depressive disorder [1]. Beyond the central nervous system, SERT serves as the primary regulator of serotonin levels in blood plasma [4]. Alterations in plasma serotonin are frequently associated with cardiovascular diseases and hemodynamic shifts.

Many common antidepressant medications exert their effects by inhibiting SERT or NET to prolong synaptic signaling [1,5]. Furthermore, DAT and NET are widely implicated in the etiology and progression of neurodegenerative diseases, most notably Alzheimer’s and Parkinson’s diseases [1,6]. These transporters also serve as the primary targets for potent drugs of abuse (such as cocaine) [7]. Despite this, the structural mechanisms driving these actions remain poorly understood, especially in the case of NET [5]. The functional architecture of MATs typically involves complex biological assemblies such as dimers or higher-order tetramers which are essential for their modulation, membrane trafficking, and turnover. Recent studies have demonstrated that the disruption of this bio-assembly can impair transporter performance, highlighting the importance of the oligomeric interface in neurotransmitter regulation [5,8].

Concurrently, the dynamic nature of transcription allows for the generation of structurally and functionally distinct proteins. Alternative splicing, initiation, and promoter usage introduce a redistribution in the coding regions of mRNA transcripts leading to the generation of multiple isoforms for a single transporter gene [9]. The functional diversity arising from the transcriptome may also impact the physiological roles of the transporters [10]. Truncated isoforms lack specific exons for the transport process but may persist in their structural similarity to the canonical transporter, especially within the transmembrane (TM) helices. Their interactions within bio-assembly were previously revealed where truncated receptors function as negative regulators when co-expressed with their full-length counterparts within the glutamate, vesicular monoamine transporter, and vesicular polyamine transporter [11–17]. Complementing this, our water soluble variant analysis established that MAT transmembrane helices exhibit structural plasticity, tolerating specific substitutions without disrupting their topology [18].

This study represents a significant expansion of knowledge on MAT bio-assembly dynamics. While the high-resolution elucidation of these interactions represents a fundamental advance in membrane protein biophysics, the pharmacological implications revealed in this study are highly substantial. Historically, the pharmacological targeting of monoamine transporters has relied almost exclusively on orthosteric or allosteric inhibitors that bind the functional transport core. In this study, we have revealed the selective inhibition of the norepinephrine transporter alongside the cross-reactivity of the specific NET-derived isoform A0A804HLI4 with both the serotonin and dopamine transporters. To resolve these interactions, we have developed a multi-level computational framework that links genomics-guided isoform identification with multimer prediction [19] and state-of-the-art molecular dynamics. Notably, we move beyond dimeric models to perform large-scale near-million-atom tetrameric simulations of the MAT canonical isoform complex and homotetramer. This unprecedented computational scale enables the high-resolution elucidation of full tetrameric assemblies and their collective dynamics within a physiologically accurate, fully atomistic lipid bilayer.

By integrating these analyses with membrane-adapted MM/PBSA binding free energy calculations for simulation trajectory, we quantitatively resolve the stability, electrostatic latches, and interface disruption patterns associated with isoform-canonical hetero-oligomers. This combinatorial approach allows us to link sequence-level isoform diversity to membrane-dependent assembly energetics, providing a thermodynamically grounded model of isoform-induced inhibition of oligomerization.

Our work highlights the impact of isoform diversity on the dimerization and tetramerization processes, providing a foundation for further studies. Finally, to contextualize these structural and thermodynamic findings, we explore the evolutionary dimensions of these heteromeric interactions. The findings suggest that truncated isoforms play a significant potential role in modulating transporter oligomerization, adding a new dimension to our understanding of neurotransmitter regulation and potentially guiding therapeutic intervention strategies. Our findings suggest that MAT oligomerization is a sensitive and pharmacologically actionable target where truncated isoforms act as endogenous titrators that shape monoaminergic signaling and disease susceptibility.

## Results and Discussions

### Descriptive Structural Bioinformatics Analyses

The serotonin, dopamine , and norepinephrine (SERT, DAT, NET) transporters, exhibits significant structural diversity through alternative splicing, generating isoforms with reduced transmembrane (TM) domain architecture. The canonical isoforms for NET (P23975; 617aa), DAT (Q01959; 620aa), and SERT (P31645; 630 aa) all maintain the standard 12-TM LeuT-fold characteristic of the solute carrier 6 family (SLC6). These canonical transporters are characterized by moderate hydrophobicity, with GRAVY indices ranging from 0.422 to 0.499, and distinct isoelectric points ranging from pI 5.89 to 7.18 (Supplementary Table 1).

The truncated isoforms display progressive TM domain loss, creating a hierarchical series of structural modules spanning from near-complete transporters to minimal membrane-anchored domains. The NET isoform P23975-3 (512aa) retains a substantial portion of the original architecture but exhibits a significant shift toward acidity (pI 6.26) and increased hydrophobicity (GRAVY 0.568). In contrast, the A0A804HLI4 isoform retains 6 TM helices (350aa). This 6-TM architecture represents a single structural half of the transporter and displays elevated basicity (pI 8.01). Further truncation is observed in the NET isoform H3BRS0 (295 aa, 5 TM), which maintains an even higher basicity of pI 8.57. The most extreme truncation, SERT isoform J3QKP3 (72aa), retains only a single transmembrane helix (1 TM) yet exhibits the highest instability index (30.93) and extreme hydrophobicity (GRAVY 1.149) (Supplementary Figures 1-18).

These systematic truncations create macromolecules that lack the complete helical bundle. For example, in the 6-TM variant A0A804HLI4, the absence of six helices likely eliminates the pseudo-symmetry necessary for independent transport function. However, the preserved transmembrane regions across these variants may allow them to function as potential regulatory partners capable of mimicking canonical interaction motifs. The distinct biophysical features of these isoforms, particularly the wide range of isoelectric points (pI 5.56 to 8.57) and varying hydrophobicities could suggest a capacity for differential environmental sorting or evolutionary pressures for these isoforms [20–23].

Phylogenetic reconstruction confirms that these isoforms represent a specialized transcriptional expansion of an ancient and conserved structural scaffold. The maximum-likelihood tree roots the MAT family with the bacterial LeuT ancestor [24] (Figure 1), demonstrating over 400 million years of structural stability across vertebrate orthologs (Mouse and Zebrafish). The close evolutionary proximity between the DAT, SERT and NET clades could suggest a mechanistic basis for the cross-reactivity.

**Figure 1.**
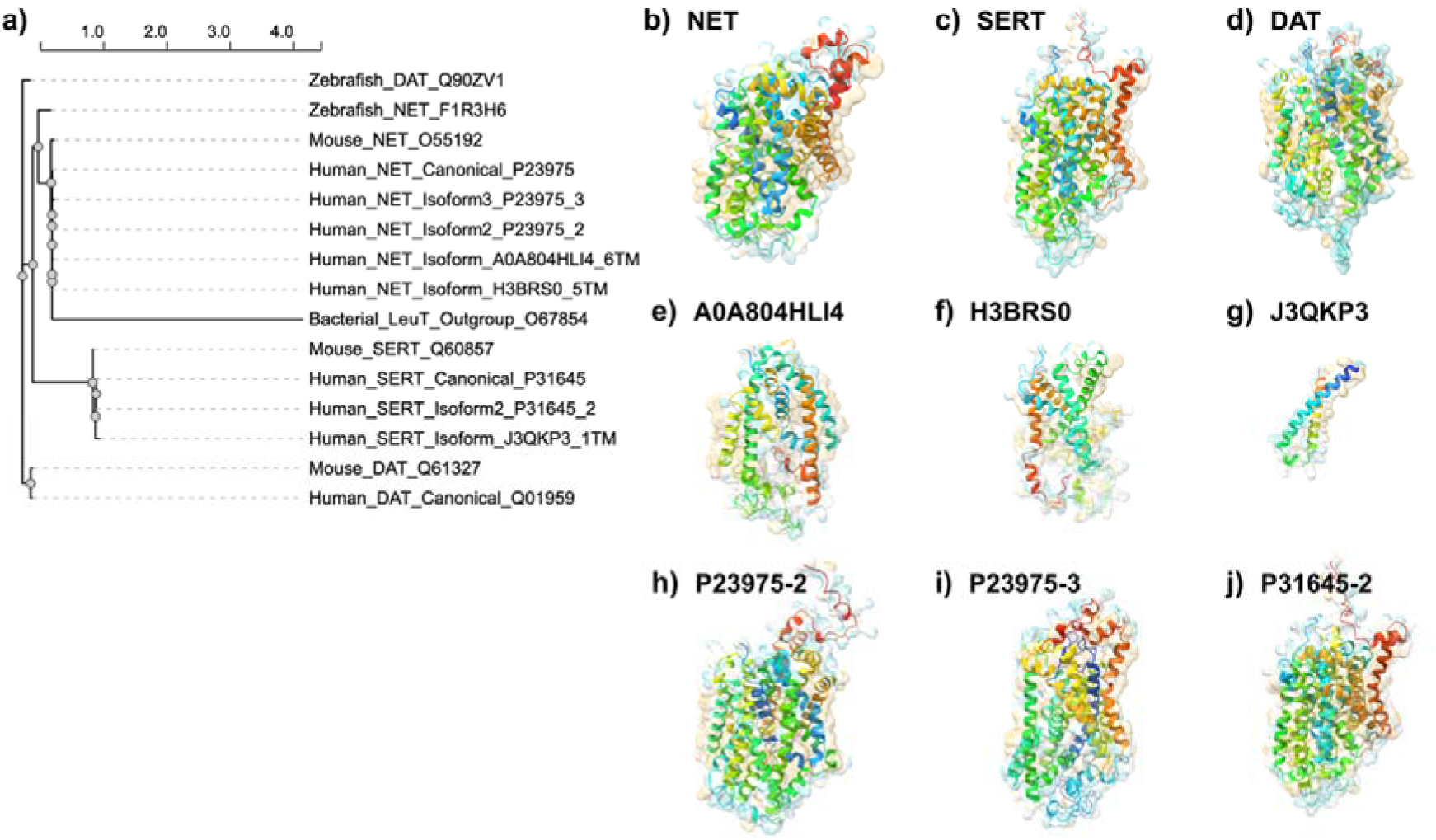
Genomic Diversity and Structural Architecture of MAT Isoforms. **(a)** Evolutionary Architecture of the SLC6 Family. Maximum-likelihood phylogenetic tree (Methods) illustrating the divergence of human monoamine transporters (SERT, DAT, NET) from the ancestral LeuT scaffold. Truncated isoforms are mapped as recent transcriptional expansions. **(b-j)** AlphaFold3 structural models of selected canonical and isoforms of human monoamine transporters (SERT, DAT, NET).

### Comparative MMGBSA Analysis

Initial MM/GBSA calculations established a hierarchical affinity profile that favored specific isoform-mediated interactions over canonical assembly in several monoamine transporter (MAT) systems. The SERT homodimer exhibited a binding free energy of -92.57 kcal/mol. Notably, this was significantly superseded by the affinity for the truncated isoform A0A804HLI4, which demonstrated a highly favorable binding energy of -143.19 kcal/mol. In the DAT system, the canonical homodimer showed substantial stability at -190.05 kcal/mol, yet the A0A804HLI4 isoform maintained strong competitive potential, reaching a comparable affinity of -179.32 kcal/mol. Similarly, while the canonical NET homodimer formed a highly stable complex (-221.75 kcal/mol), its interaction with the A0A804HLI4 variant remained energetically significant at -125.83 kcal/mol. In contrast, complexes involving the H3BRS0 variant yielded weaker affinities across all three transporters, ranging from -36.36 to -63.43 kcal/mol (Table 1). These preliminary static results underscore a potent decoy potential specifically for the A0A804HLI4 truncated variant, which was subsequently investigated through dynamic full-atom simulations.

**Table 1.** Interface composition and binding free energies (ΔΔGs) of the sampled homodimer and NET truncated isoform heterodimer complexes.

| Receptor | Ligand<sup>1</sup> | Contact Composition<sup>2</sup> | | Binding free energy ($\Delta\Delta G$)(kcal/mol)<sup>3</sup> |
| --- | --- | --- | --- | --- |
|  |  | (Receptor) | (Ligand) |  |
| **DAT** | Homodimer (Canonical) | PHE171, GLU174, TRP238, PHE168 | PHE171, THR173, TRP238, GLU174 | -190.05 |
|  | A0A804HLI4 | TRP238, PHE168, LEU246, PHE253 | TRP235, PHE164, LEU239, LYS254 | -179.32 |
|  | H3BRS0 | LYS65, LYS264, ARG283, THR286 | LEU169, LYS61, THR258, SER259 | -36.36 |
| **NET** | Homodimer (Canonical) | ARG587, ARG49, TRP553, PHE435 | ARG49, LEU45, TRP553, PHE435 | -221.75 |
|  | A0A804HLI4 | PHE167, PHE164, LEU169, TRP235 | PHE167, TRP235, PHE164, LYS261 | -125.83 |
|  | H3BRS0 | LEU169, THR283, PHE403, THR402 | PHE284, TYR287, PHE167, LEU169 | -53.91 |
| **SERT** | Homodimer (Canonical) | PHE422, PHE191, ASP193, THR421 | PHE422, ASP193, ALA419, PRO418 | -92.57 |
|  | A0A804HLI4 | ASP193, PHE191, ILE188, PHE422 | PHE164, PHE167, TRP235, LEU169 | -143.19 |
|  | H3BRS0 | PHE422, PHE287, ILE425, THR421 | ASN89, ILE104, ILE96, LEU100 | -63.43 |

<sup>1</sup>The Uniprot Entry ID of the isoform (ligand of the complex).
<sup>2</sup>Interface composition of the complex (highest energy contributing residues).
<sup>3</sup>Residue binding free energies are calculated according to MMGBSA algorithm.

Although MM/GBSA provides an end-point screening method to rank interaction potentials, it utilizes an implicit continuum solvent model and lacks conformational entropy penalties, which inherently overestimates the absolute magnitude of binding free energies [25]. To determine whether these truncated variants act as true thermodynamic decoys in a biologically accurate environment, it was imperative to subject these static predictions to dynamic, explicit-solvent validation.

### Comparative Energetics and Thermodynamic Affinity of MAT Complexes

To evaluate the stability of these interactions, the complexes were analyzed through dynamic full-atom simulations embedded in an explicit lipid bilayer environment. In this membrane context, binding free energies are naturally tempered by structural relaxation, thermal fluctuations, and explicit lipid and solvent screening. Under these realistic conditions, the assessment of binding free energy profiles revealed critical differentials between canonical MAT homodimers and the isoform-coupled complexes (Figure 2).

**Figure 2.**
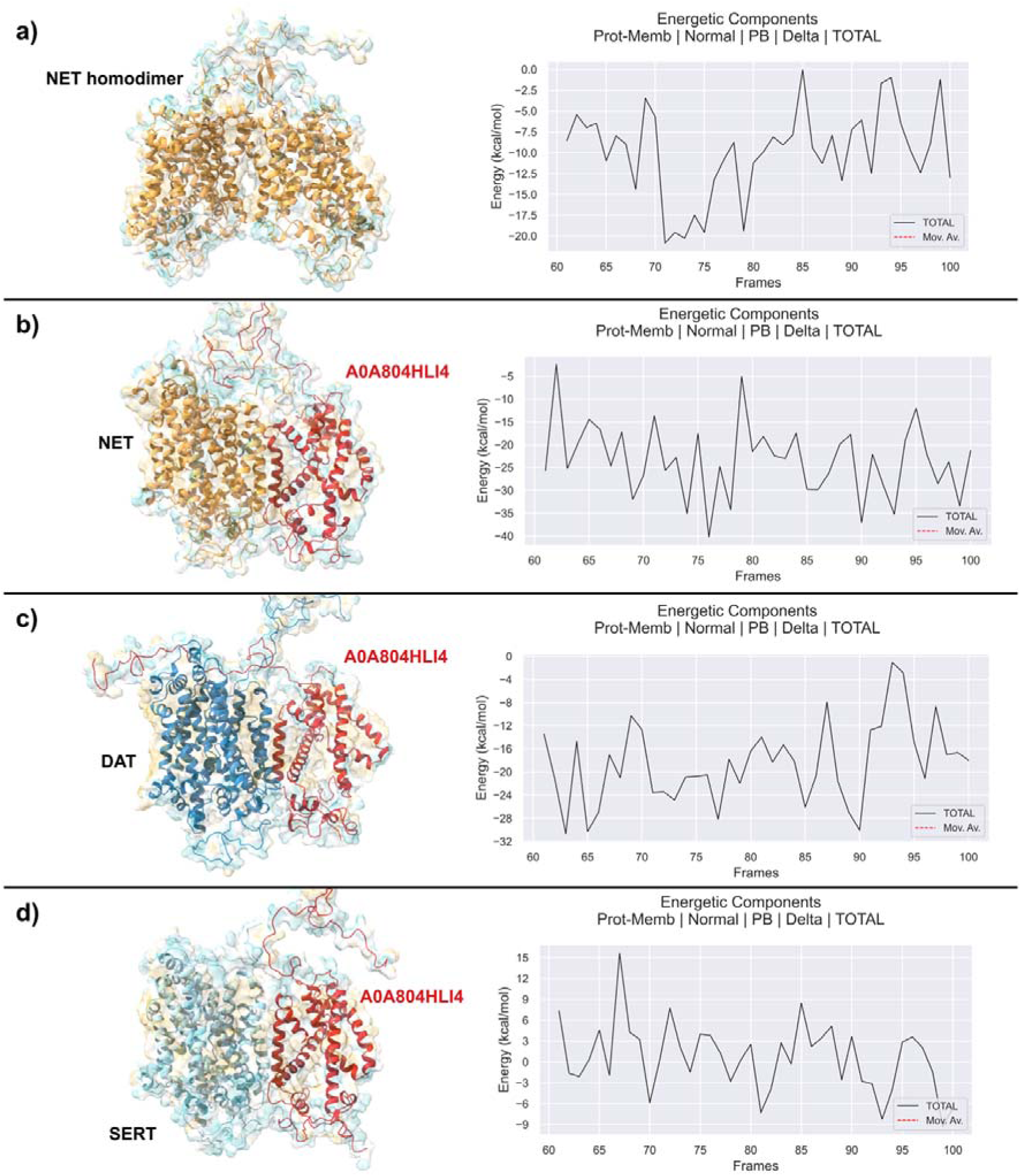
Comparative binding free energy profiles of MAT homodimeric and isoform-heterodimeric complexes. Binding energetics and thermodynamic preference of the A0A804HLI4 truncated variant across MAT family members. **(a)** NET homodimer control trajectory exhibiting a baseline binding energy fluctuating near neutral values. **(b)** NET-A0A804HLI4 and **(c)** DAT-A0A804HLI4 heterodimers demonstrate significant exergonic profiles, frequently stabilizing at negative values below -35 kcal/mol and -28 kcal/mol, respectively. **(d)** SERT-A0A804HLI4 dimer exhibits a comparatively higher free energy profile than NET or DAT dimers, indicating a relative thermodynamic preference for NET and DAT templates in the dimeric state.

While the canonical DAT-DAT homodimer exhibited a fluctuating energetic baseline (averaging between -3 and +9 kcal/mol, indicative of a transient association) the heterodimer involving the truncated variant A0A804HLI4 (6 TMs) demonstrated a stable exergonic profile. The DAT-A0A804HLI4 trajectory stabilized strictly at negative values, frequently reaching below -28 kcal/mol. This magnitude of energy in an explicit, membrane-bound dynamic system represents a remarkably tight protein-protein interaction. Similarly, the NET-A0A804HLI4 complex exhibited consistent stabilization with values dipping below -35 kcal/mol.

In contrast, the SERT-A0A804HLI4 dimer demonstrated comparatively lower binding-energy indicating that the A0A804HLI4 isoform maintains a thermodynamic preference for NET and DAT over SERT in a dimeric configuration. This energetic disparity suggests that truncated variants possess optimized interfaces capable of outcompeting native oligomerization. The thermodynamic profiles observed across MAT-A0A804HLI4 complexes characterize a state of high thermodynamic stability compared to canonical homodimers, which often display higher variance and more positive baselines.

The observed gradients suggests that the truncated isoform acts as a potential selective modulator for monomers. This establishes a thermodynamic hierarchy where the isoform-bound complexes are favored over functional homodimers, particularly in the case of NET and DAT. This preference likely dictates the availability of functional transporters at the membrane by sequestering monomers into heterodimeric states.

### General Modulatory Mechanism via Cross-Reactivity

Beyond parental interactions, these results expose a layer of regulation via inter-transporter cross-talk. Potential evidence of cross-reactivity was revealed in simulations involving the NET-derived isoform A0A804HLI4. When evaluated against non-parental transporters, this isoform demonstrated a high capacity for promiscuous binding. Both the DAT-A0A804HLI4 and NET-A0A804HLI4 complexes maintained stable binding profiles that outperformed the baseline stability of native assemblies. This could suggest a competitive interference mechanism where a single truncated variant acts as a general potential inhibitor across the SLC6 family. The energetic trajectories indicate that the isoform effectively interferes with the assembly of its own parent transporter while simultaneously destabilizing serotonin and dopamine transporter oligomerization. Such cross-reactivity implies that the dysregulation of a single isoform could have systemic impacts on multiple monoaminergic pathways. Furthermore, the stability of these cross-reactive complexes may suggest that the isoform interface is conserved or sufficiently flexible to accommodate the structural nuances of different MAT family members.

### Physicochemical Drivers of Isoform Interference

The distinct physicochemical properties of the truncated isoforms emerge as potential key drivers of their regulatory interference. From an evolutionary perspective, these extreme physicochemical shifts (particularly the transition toward highly basic and altered hydropathy profiles within the membrane) are likely not stochastic. The stability of such divergent transmembrane architectures requires precise compositional balancing. As we previously demonstrated in a large-scale analysis of human alpha-helical membrane proteins, substitutions between nonpolar and polar residues may carry high pathological penalties; however, functional diversification can be achieved when these shifts are mitigated by specific co-evolutionary residue pairing and intermediate substitution pathways [26]. Applied to the MAT family, the marked divergence in isoelectric points (pI) may suggest a selected compositional bias that provides a theoretical basis for electrostatic attraction and charge complementarity. For instance, the basic nature of the truncated variant A0A804HLI4 (pI 8.01) provides an electrostatic contrast to the slightly acidic or near-neutral canonical DAT (pI 6.46) and NET (pI 7.18) architectures. This elevated basicity may dictate targeted interactions with negatively charged membrane phospholipids, anchoring the isoforms to specific regions of the inner leaflet. Coupled with varying hydropathy profiles, this complementary charge distribution could not only thermodynamically favor heterodimeric assembly but also facilitate selective phase partitioning into ordered lipid microdomains [22].

Notably, this targeted electrostatic shift is not exclusive to truncated variants (Supplementary Table 1). Interestingly, the NET isoform P23975-2, retains the 12-TM architecture, similar mass, and baseline hydrophobicity of the canonical transporter, yet exhibits a dramatic shift toward basicity (from pI 7.18 to 8.44). At physiological pH, this elevated basicity dictates a substantial increase in the positive charge of the transporter. Such physicochemical alteration, achieved without compromising the core hydrophobicity, likely enhances the isoform’s electrostatic affinity for anionic phospholipid headgroups at the inner leaflet of the membrane. Consequently, this targeted isoelectric shift may drive the differential lateral segregation of the isoform into specific lipid microdomains or alter its interactome with intracellular scaffolding proteins. While exploring these interactions is beyond the immediate scope of this study, thus, future investigations are necessary to definitively resolve the precise structural and physiological contributions of this isoelectric shift within a cellular context.

### Thermodynamic Architecture of Near-Million-Atom SERT Tetrameric Complexes

Near-million-atom tetrameric simulations provided additional insights into the kinetic stability and mechanism of these isoform-induced disruptions. The progression from dimeric affinity to high-order tetrameric architectures reveals a thermodynamic lock within the canonical serotonin transporter (SERT) homotetramer that is absent in isoform-coupled systems (Figure 3). Analysis of the <u>T</u>otal <u>D</u>ecomposition <u>C</u>ontribution (TDC) over the final trajectory frames shows stabilizing interactions in the homotetramer. Primary adjacent interfaces maintain a consistent energetic baseline with total binding free energies consistently favoring a stable state.

**Figure 3.**
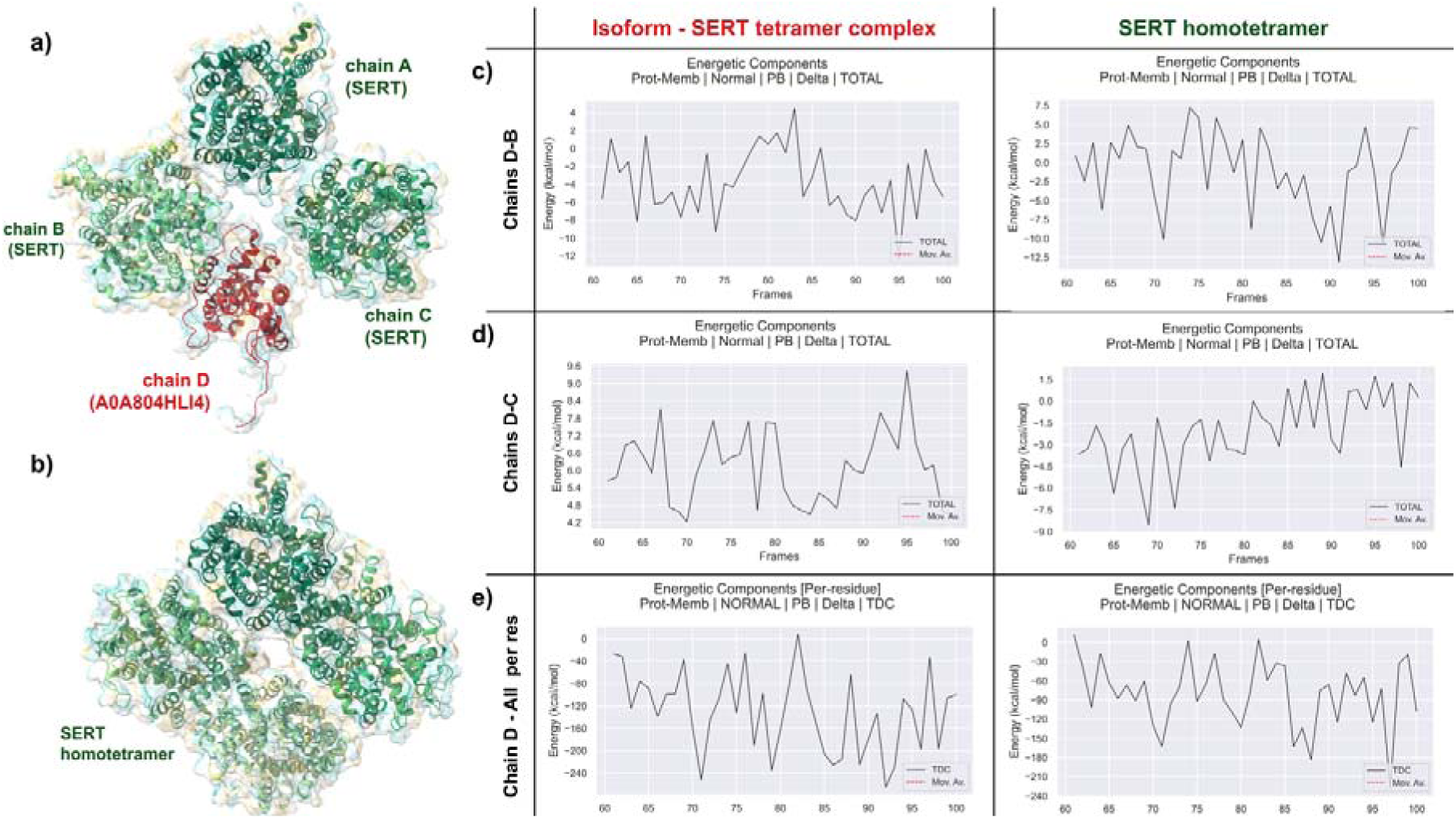
Quaternary assembly disruption of the SERT tetramer by the A0A804HLI4 truncated isoform. Thermodynamic destabilization and interfacial repulsion in hybrid SERT tetrameric assemblies. **(a)** Structural model of the hybrid SERT tetramer (A3B stoichiometry) with the A0A804HLI4 isoform (Chain D) highlighted in red. **(b)** Canonical SERT homotetramer (A4) showing symmetric quaternary architecture. **(c)** Interaction energy for the Chain D-Chain B interface remains negative, serving as the primary contact site for the variant. **(d)** The Chain D-Chain C interface exhibits a consistent repulsive shift, with total binding energy remaining positive throughout the trajectory (range: +4.2 to +9.4 kcal/mol). **(e)** Total Decomposition Contribution (TDC) line plots for Chain D in the hybrid complex versus the homotetramer. The transition to positive energetics at the D-C interface indicates that the isoform architecture is fundamentally incompatible with canonical assembly motifs, leading to global allosteric destabilization. SERT N-terminal loops are removed for clarity.

While canonical tetramers exhibited symmetric conformational motions, assemblies containing isoform variant showed persistent structural asymmetry. The introduction of the A0A804HLI4 truncated isoform at the Chain D position triggers a structural destabilization of this architecture. While the canonical system maintains a stabilizing baseline at the D-C interface, the isoform system exhibits consistent repulsion at this site. The total binding energy for the Chain D (Isoform)-Chain C interface remains entirely positive throughout the analyzed trajectory, fluctuating between +4.2 kcal/mol and +9.4 kcal/mol. Once the isoform bound to a canonical protomer the interface remained locked throughout the evaluation period.

Significantly, the Chain D (Isoform)-Chain B interface emerges as the primary contact site for the variant, exhibiting a negative affinity that acts as a competitive inhibitor of proper quaternary assembly. The TDC line plots for the isoform D-B interface show localized energy dips reaching as low as -12 kcal/mol in total energy. However, this localized affinity at the D-B site appears to be a non-native compensatory mechanism for the failure of the D-C interface. By anchoring to Chain B while simultaneously repelling Chain C, the isoform creates an unbalanced structural torque that prevents the tetramer from reaching a global equilibrium.

Despite local frustrations at the tertiary level, the overall enthalpic gain was sufficient to maintain complex integrity. This kinetic persistence suggests that isoform-canonical interactions represent stable regulatory states rather than transient associations. By locking the tetramer into a non-symmetric state, the isoform not only reduces the number of functional subunits but also disrupts the mechanical cooperation of the remaining WT protomers.

### Residue-Level Energetics of Dimeric Assembly

To resolve the atomic-level drivers of energetic divergence within the isolated dimeric complexes, <u>P</u>er-<u>R</u>esidue <u>E</u>nergy <u>D</u>ecomposition (PRED) was employed (Figure 4). This analysis indicated a mechanistic shift between the distributed hydrophobic packing of canonical forms and the highly specific residue interactions found in isoform dimer complexes. The selectivity of the isoform for NET appears to be driven by Gln236, which functions as a specific molecular latch; it exhibits a favorable energetic contribution in the NET-isoform dimer that is absent in the DAT complex.

**Figure 4.**
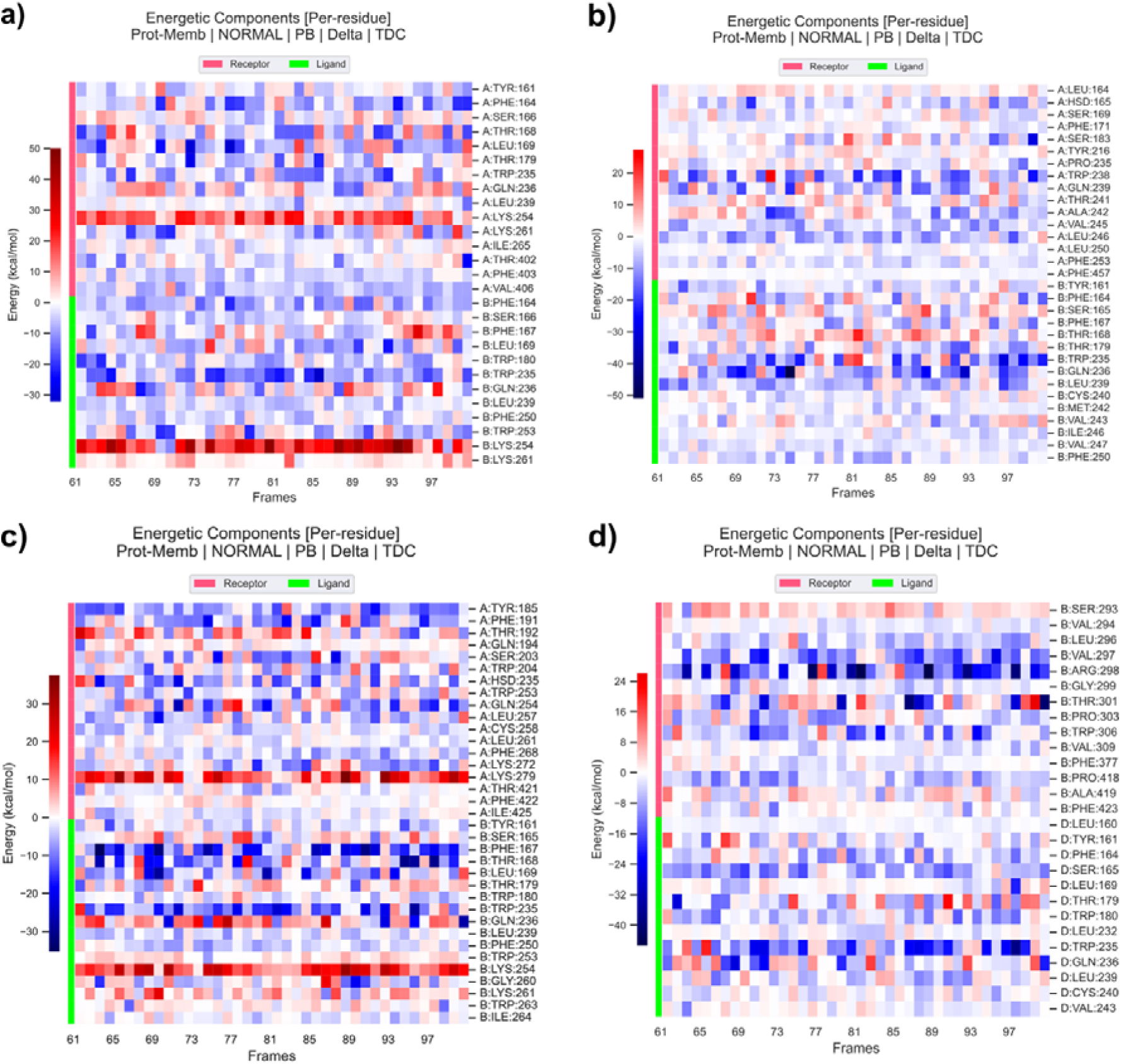
Atomistic drivers of MAT-isoform selectivity and allosteric interference. High-resolution heatmaps illustrate the time-resolved Total Decomposition Contribution (TDC) for key residues at the isoform-transporter interface. **(a)** NET - A0A804HLI4 isoform heatmap. **(b)** DAT - A0A804HLI4 isoform heatmap. **(c)** SERT - A0A804HLI4 isoform dimer and **(d)** SERT - A0A804HLI4 isoform tetramer heatmaps. In the tetrameric state, residues Ser165 and Trp235 exhibit enhanced stabilization (deeper blue) relative to the dimeric state.

**Figure 5.**
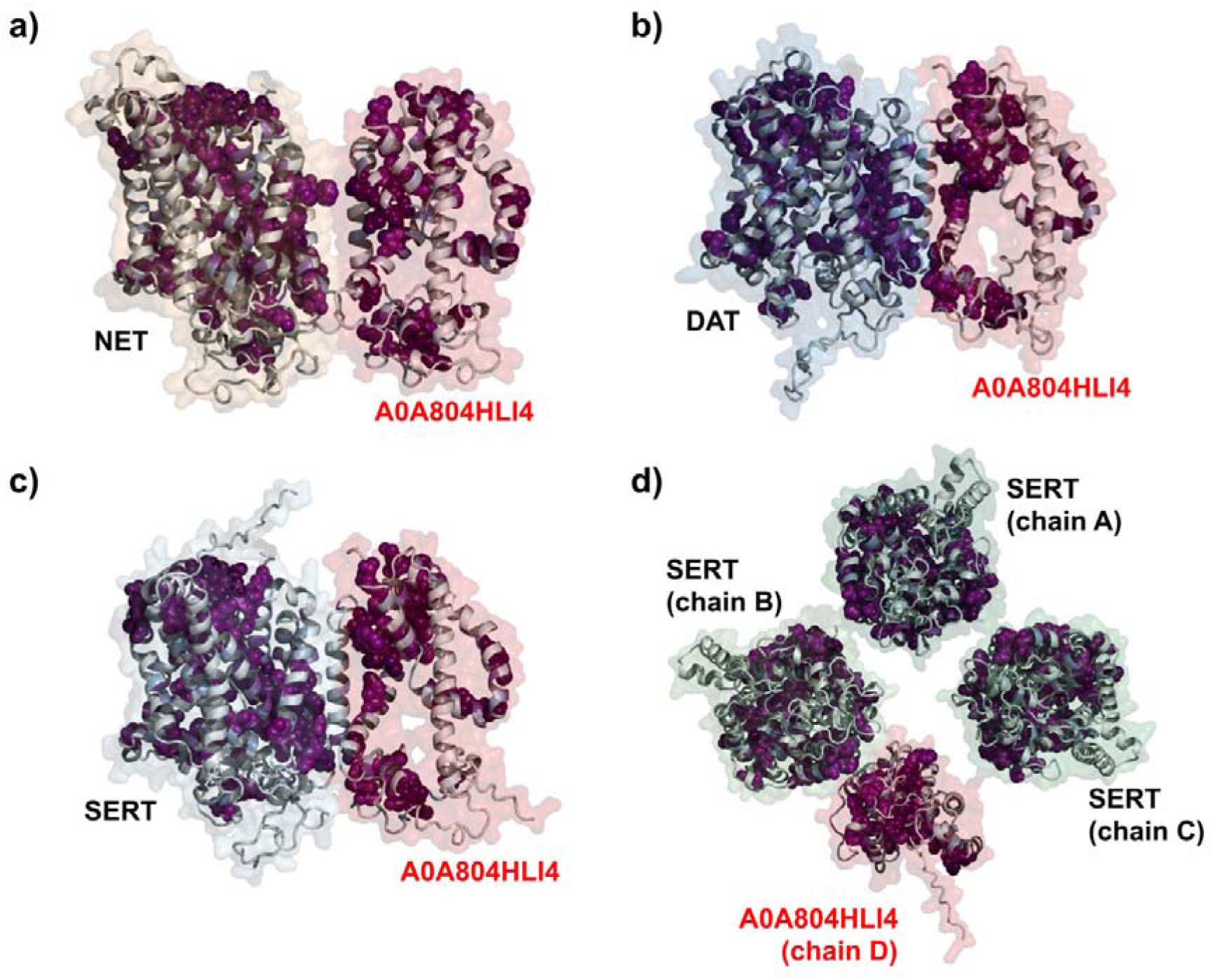
Evolutionary coupling analysis of hybrid monoamine transporter complexes. Structural representation of predicted evolutionary couplings enrichments (purple spheres) mapped onto the models of monoamine transporter (MAT) family complexes (large N-terminal loops are removed for clarity). **(a)** NET-A0A804HLI4 heterodimer, **(b)** DAT-A0A804HLI4 heterodimer, and **(c)** SERT-A0A804HLI4 heterodimer. **(d)** SERT hybrid tetramer (A3B stoichiometry) illustrating the distribution of coupling constraints across the quaternary assembly.

Across all MAT family interactions, Trp235 (A0A804HLI4) emerged as a universal hydrophobic anchor, providing a dominant stabilization point for isoform association in the dimeric state. In canonical homodimers, stabilization is mediated primarily by a distributed network of aromatic clusters, which lack the localized intensity observed in isoform-coupled systems. The identification of these specific hotspots suggests that the dimeric isoform-transporter interface is not only a byproduct of random hydrophobic collapse but a coordinated structural interaction. Interestingly, the deep energetic wells provided by Trp235 ensure a baseline affinity, while residues like Gln236 (especially for DAT interactions) (Figure 4b) provide the necessary specificity for selective transporter targeting.

### Residue-Level Structural Destabilization of the Tetrameric Architecture

Expanding this atomistic analysis from the isolated dimer to the larger quaternary architecture, high-resolution per-residue heatmaps reveal the potential drivers behind the structural failure of the tetramer and the compensatory stabilization of the D-B primary contact site. In the canonical tetrameric assembly, stability is maintained by a sophisticated residue-level network. In contrast, the isoform-disrupted D-C interface exhibits a loss of these stabilizing motifs, which are replaced by an energetic penalty. This repulsion implies that the truncated variant exposes structural regions that sterically clash with neighboring canonical subunits within the tetramer.

Heatmap analysis reveals that residue Ser165 become significantly more favorable in the SERT tetramer than in the isolated SERT dimer (Figure 4b). This energetic strengthening of specific hotspots suggests that the isoform is forced into a tighter packing arrangement with Chain B to offset the repulsion it experiences from Chain C. While these residues provide strong anchoring points, they exist in a state of structural misfit, forcing the protein backbone into strained conformations.

Finally, the allosteric ripple effect of the isoform is visible at distal interfaces composed purely of canonical proteins, indicating that the disruption is a global tetrameric event. Distal interfaces exhibit new repulsion bands in the presence of the isoform, shifting previously stabilizing residues to a destabilizing role. Energy decompositions demonstrate that this shift toward higher free-energy is driven by a dominant solvation penalty that overcomes the reduced van der Waals stabilization. These findings characterize A0A804HLI4 as a structural variant that likely utilizes both localized high-affinity binding and distal allosteric strain to prevent the formation of functional SERT tetramers.

### Interface evolutionary coupling of MAT complexes

To evaluate whether the thermodynamic hyper-affinity observed in our MD simulations reflects biologically relevant interactions rather than stochastic packing, molecular dynamic simulations was integrated with evolutionary enrichment analysis (dynamics-aware evolutionary profiling) [27]. By mapping co-evolutionary scores onto the physical contact residues identified in the DAT (Q01959) and NET (P23975) complexes with the A0A804HLI4 isoform, a high convergence between structural stability and evolutionary selection was observed. This convergence defines a set of hypothetical evolutionary hotspots that anchor the isoform-canonical interface. In the DAT-A0A804HLI4 complex, the interaction is dominated by a remarkable evolutionary anchor at Gln239 (DAT) and Gln236 (Isoform). Our decomposition analysis identified this residue pair as the primary structural stabilizer, exhibiting a substantial interaction energy (IE) of -28.46 kcal/mol and -24.08 kcal/mol, respectively. Crucially, these positions also exhibit high enrichment scores (5.06 for DAT; 3.67 for Isoform), suggesting that the specific chemistry of this glutamine-mediated contact is maintained under significant selective pressure. This “glutamine-latch” is further supported by Leu164 (IE: -9.08 kcal/mol; Enrichment: 3.86) and Tyr216 (IE:-9.00 kcal/mol; Enrichment: 2.68) on the DAT side, forming a co-evolved hydrophobic-polar network that likely facilitates the isoform’s competitive binding.

A similar pattern of evolutionary coupling emerged in the NET-A0A804HLI4 interface. The residue Tyr161 acts as a critical structural and evolutionary node. In the A0A804HLI4 isoform, Tyr161 displays an exceptionally high enrichment score of 7.78, corresponding to a strongly favorable interaction energy of -5.72 kcal/mol in the NET-coupled state. Its partner in the canonical NET transporter, Tyr161, similarly exhibits high enrichment (5.44) and substantial stabilization (IE: -9.27 kcal/mol). The presence of such high evolutionary signatures at the exact coordinates of maximum physical interaction implies that these truncated isoforms have retained, or perhaps optimized, specific interaction fingerprints that allow them to effectively intercept canonical transporters. Mapping these scores indicates that the affinity of the A0A804HLI4 variant relies heavily on several highly conserved residues. For instance, in the NET complex, Gln236 and Val406 show enrichment scores of 4.29 and 7.54, respectively, marking them as evolutionarily primed for interface formation.

This convergence of MD-derived energetics and sequence-based evolutionary metrics suggests that the regulatory/inhibitory role of truncated MAT isoforms is encoded within their primary sequence. The high enrichment scores at the interface imply that these residues are under significant evolutionary constraint, likely because they are essential for the formation of the isoform-canonical heterodimer. These findings could reinforce the decoy hypothesis, characterizing these isoforms as evolutionarily conserved structural terminators potentially designed to titrate MAT oligomerization through interface coupling.

### Conclusion and Potential Implications

The integrated computational framework presented here reveals that monoamine transporter regulation extends far beyond simple genomic expression and involves a sophisticated layer of quaternary sabotage mediated by the A0A804HLI4 isoform. By transitioning from traditional monomeric models to the near-million-atom scale of tetrameric assembly, we have exposed a potential endogenous mechanism where alternative splicing operates as a precision titrator of synaptic signaling. While our atomistic lipid bilayer represents a significant step toward physiological realism, it is important to acknowledge that this model does not yet capture the full heterogeneity of the presynaptic membrane, such as diverse cholesterol gradients or auxiliary scaffolding proteins that likely modulate these interactions in a living cell.

Interestingly, the observed cross-reactivity between the NET-derived A0A804HLI4 variant and the broader MAT family introduces a shift in our understanding of neurotransmitter cross-talk. While previous studies on vesicular transporters and glutamate systems suggested localized interference, our data implies that a single splice variant can exert systemic influence across multiple monoaminergic pathways. This promiscuous binding capacity may provide a structural explanation for the interconnected nature of neuropsychiatric comorbidities where transcriptional dysregulation in one transporter locus manifests as a multi-systemic failure of neurochemical homeostasis. Although our MMPBSA data identifies stable interaction nodes, the absolute affinities in vivo may be further influenced by the local concentration of transporters and the kinetic competition for trafficking chaperones.

Our near-million-atom simulations provide a unique perspective on the macro-structural collapse of these complexes through the poison subunit effect. This mechanism demonstrates that the inclusion of a single truncated protomer creates an unbalanced structural torque that ripples across the entire tetramer, forcing distal interfaces into high-free-energy states. This allosteric destabilization effectively renders the million-atom assembly transport-incompetent, representing a rapid post-translational method to downregulate synaptic reuptake. We must consider, however, that the hundreds of nanoseconds of MD trajectory, while massive in scale, may still miss rarer conformational transitions or the slow kinetics of higher-order dissociation. Future efforts integrating these findings with in vivo single-molecule FRET might be essential to capture the real-time dissociation kinetics of these heteromeric complexes.

The convergence of atomistic interaction energies with evolutionary enrichment scores additionally confirms that the A0A804HLI4 interface is a functionally conserved regulatory site. The high selection pressure on residues such as Tyr161 and Gln236 indicates that these interference nodes are evolutionarily primed to facilitate high-fidelity steering of the isoform toward its target. This structural mimicry allows the truncated variant to outcompete native subunits with an irreversible energetic preference, acting as a regulatory switch to manage the ratio of active-to-inactive transporters. While AlphaFold3 provides unprecedented structural accuracy, the inherent flexibility of truncated variants suggests they may exist in multiple metastable states that require further characterization via cryo-electron microscopy or NMR spectroscopy to fully resolve the MAT proteome.

From a clinical standpoint, these non-canonical interfaces offer a novel template for the next-generation of precision therapies. Therapeutic intervention could focus on decoy-capping peptidomimetics or hydrocarbon-stapled alpha-helical peptides designed to neutralize the isoform’s evolutionary hotspots and rescue functional monomers from sequestration. Alternatively, molecular glues could be engineered to reinforce the native hydrophobic clefts of canonical dimers to render the functional assembly against isoform-mediated disruption. Ultimately, recognizing the principle of isoform-mediated quaternary sabotage allows for the development of targeted strategies to restore the structural integrity and functional capacity of the monoamine transporter system in diseased states.

## Methods

### Protein Sequences and Physicochemical Characterization

Primary protein sequences for the canonical human monoamine transporters, including the norepinephrine transporter (NET, P23975), dopamine transporter (DAT, Q01959), and serotonin transporter (SERT, P31645), were accessed from UniProtKB [28]. To explore the structural diversity generated by alternative splicing, a genomic screening was conducted using the Ensembl genome browser and UniProt mapping tools to identify all protein-coding isoforms [28, 29]. The selection criteria focused on isoforms with sequence lengths > 10% and < 90% of the full-length canonical proteins to ensure structural relevance for membrane-embedded simulations (P23975-2 and P31645-2 have also been added to the tables for comparison purposes). Membrane topology and transmembrane (TM) helix architecture were visualized and cross-validated using the Protter server [30]. Physicochemical properties, including molecular weight, and Grand Average of Hydropathy (GRAVY) scores and Kyle-Doolittle analysis, were calculated via the Expasy ProtParam tool [31–33] and our Evolutionary Statistics Toolkit (version 1.2.2) [34]. Hydrophilic variants exhibiting negative GRAVY scores were excluded to focus on membrane-integrated modules. Following this initial characterization, the NET-derived 6-TM variant A0A804HLI4 was identified as a primary regulatory candidate.

### Phylogenetic Reconstruction and Evolutionary Sampling

To evaluate the evolutionary divergence of the monoamine transporter (MAT) family, canonical sequences for human SERT (SLC6A4), DAT (SLC6A3), and NET (SLC6A2) were aligned with their respective orthologs from *Mus musculus* (Mouse) and *Danio rerio* (Zebrafish). The bacterial leucine transporter (LeuT), from *Aquifex aeolicus* (O67854), was utilized as the structural outgroup to root the phylogeny [28,29]. Phylogenetic analysis was conducted using the NGPhylogeny.fr default workflow [35,36]. Multiple sequence alignment was performed via MAFFT (flavour: auto; gap opening penalty: 1.53; gap extension penalty: 0.123) [37]. The alignment was refined using BMGE (sliding window: 3; entropy threshold: 0.5) to remove ambiguous regions [38]. Phylogenetic inference was performed using PhyML+SMS (Smart Model Selection) with the LG substitution model and ML equilibrium frequencies. Tree topology was optimized via Subtree Pruning and Regrafting (SPR), with the proportion of invariant sites and gamma distribution parameters estimated from the data. Initial tree construction was supplemented by FastME using balanced minimum evolution (BalME) [39,40].

### AlphaFold3 Predictions and Systematic Model Ranking

Structural models for all identified isoform-canonical monomers, dimers, and tetrameric complexes were generated using the AlphaFold3 server [19] to produce high-confidence starting configurations. To identify the most potent regulatory partners, a hierarchical ranking strategy was employed across the isoform library. Preliminary screening was conducted using the Molecular Mechanics with Generalized Born and Surface Area method implemented via the HawkDock server [41–42]. All models were pre-processed using the tleap module in Amber16 to add missing hydrogens and heavy atoms. The ranking process utilized a variable dielectric generalized Born model to evaluate the competitive binding energetics of each isoform against its native canonical homodimer. This systematic evaluation identified A0A804HLI4 as exhibiting superior competitive affinity, justifying its integration into the near-million-atom tetramer simulations.

### Near-Million-Atom Plasma Membrane Systems

To investigate the macro-structural impact of isoform integration, all-atom tetrameric membrane systems were constructed using the CHARMM-GUI Membrane Builder [44–46], similar to our previous studies [47,48]. Protein spatial alignment within the lipid bilayer was optimized using the PPM 2.0 algorithm [49], which accounted for the anisotropic dielectric properties of the membrane-water interface. The complex lipid environment was designed to mimic a representative human plasma membrane, consisting of cholesterol (25%), POPC (30%), POPE (20%), POPS (5%), POPI (5%), and palmitoyl-sphingomyelin (PSM, 15%). These extensive assemblies reached a near-million-atom scale (approximately 950,000 atoms) within the simulation box. System neutrality and a physiological ionic strength of 0.15 M were achieved through the addition of K+ and Cl- ions. The CHARMM36m all-atom force field [50] was applied to all system components, including protein, lipids, and ions.

### Molecular Dynamics Simulations

Molecular dynamics simulations were executed using GROMACS 2024.3 [51] on AlphaFold-predicted structures embedded in the lipid bilayer. High-performance computational analysis was performed on a Google Colab cluster equipped with NVIDIA L4 GPUs and Intel® Xeon® CPUs. To maximize throughput for the million-atom systems, the GROMACS software was recompiled with CUDA support for enhanced GPU parallelization [52]. Energy minimization was performed using the descent algorithm with a force tolerance of 1000 kJ/mol/nm. System equilibration followed a rigorous six-step protocol involving the gradual reduction of position restraints on protein heavy atoms and lipid headgroups. Production MD simulations were carried out for 50 ns in the NPT ensemble at 310 K and 1 bar. Electrostatic interactions were computed via the Particle Mesh Ewald method with a 1.2 nm cutoff, and all bonds containing hydrogen atoms were constrained using the LINCS algorithm to facilitate a 2-fs timestep [53].

### Binding Free Energy and Decomposition Analysis

Binding free energies for both the dimeric screening set and the production tetrameric trajectories were computed using the Molecular-Mechanics Poisson-Boltzmann Surface-Area method via the gmx_MMPBSA tool [54,55]. Analysis focused on the stabilized, equilibrated portions of the trajectories, employing a three-dielectric model where the membrane, solute, and solvent were assigned dielectric constants of 7.0, 4.0, and 80.0, respectively. Total electrostatic energies were calculated using the particle-particle particle-mesh (P3M) method. To resolve the structural basis of interface failure and hyper-affinity, per-residue energy decomposition was performed to calculate the Total Decomposition Contribution (TDC) for each residue. This allowed for the isolation of specific polar and aromatic clusters contributing to the total binding affinity. Standard error of the mean (SEM) was calculated through uncertainty propagation, and interface contacts within 4 Å were analyzed and visualized using UCSF ChimeraX [56].

### Co-evolutionary Profiling and Hotspot Mapping

Evolutionary couplings (ECs) were determined via the EVcouplings [57] server using a maximum entropy model constrained by multiple sequence alignment (MSA) statistics. Length-normalized bitscores were utilized to compare the sequence conservation of canonical MATs with their truncated isoforms. The resulting enrichment scores were integrated with the atomistic interaction energies (IE) derived from the MD simulations, employing a dynamics-aware evolutionary profiling approach [27]. This integrated mapping allowed for the identification of evolutionary hotspot residues that demonstrate both high selective pressure and significant structural stabilization.

## Supplementary information

Supplementary_Information.pdf

## Data availability statement

The data and custom code supporting this article are freely available to the scientific community, in our public GitHub repository at https://github.com/karagol-taner/net-dat-sert-truncation-cross-reactivity. Any additional thermodynamic or other data not included in the main text or supplementary materials are available from the corresponding authors upon reasonable request.

## Ethics approval

Ethics approval was not required for this computational study as it did not involve animal subjects, human participants, and identifiable data.

## Consent to participate

Not applicable. This computational study did not involve human participants.

## Consent for publication

Not applicable. This computational study did not involve human participants.

## Author contributions

A.K and T.K contributed equally to this work.

