## Supplementary_Information.pdf for "Serotonin, dopamine, and norepinephrine transporter assembly is selectively disrupted by a NET truncation isoform as revealed through near-million-atom simulations"

Taner Karagöl[1,¶,*], Alper Karagöl[1,¶,*],
[1]Istanbul Faculty of Medicine, Istanbul University, Istanbul, Turkey

[¶]These authors contribute equally.
*To whom the correspondence should be addressed.

Taner Karagöl,
ORCID: 0009-0005-1011-7661

Alper Karagöl,
ORCID: 0009-0001-7864-0732

### Table of Contents

1. Table S1. Computationally mapped and predicted potential truncated isoforms of vesicular amine transporter subfamily (VATs). Pg. 3
2. Figure S1. Sequence and topology visualization of the canonical NET isoform P23975 from Protter. Pg. 4
3. Figure S2. Sequence and topology visualization of the NET isoform P23975-2 from Protter. Pg. 5
4. Figure S3. Sequence and topology visualization of the NET isoform P23975-3 from Protter. Pg. 6
5. Figure S4. Sequence and topology visualization of the NET isoform A0A804HLI4 from Protter. Pg. 7
6. Figure S5. Sequence and topology visualization of the NET isoform H3BRS0 from Protter. Pg. 8
7. Figure S6. Sequence and topology visualization of the canonical DAT isoform Q01959 from Protter. Pg. 9
8. Figure S7. Sequence and topology visualization of the canonical SERT isoform P31645 from Protter. Pg. 10
9. Figure S8. Sequence and topology visualization of the SERT isoform P31645-2 from Protter. Pg. 11
10. Figure S9. Sequence and topology visualization of the SERT isoform J3QKP3 from Protter. Pg. 12
11. Figure S10. Kyte-Doolittle hydropathy plot of the canonical NET isoform P23975. Pg.13
12. Figure S11. Kyte-Doolittle hydropathy plot of the NET isoform P23975-2. Pg. 14
13. Figure S12. Kyte-Doolittle hydropathy plot of the NET isoform P23975-3. Pg. 15
14. Figure S13. Kyte-Doolittle hydropathy plot of the NET isoform A0A804HLI4. Pg. 16
15. Figure S14. Kyte-Doolittle hydropathy plot of the NET isoform H3BRS0. Pg. 17
16. Figure S15. Kyte-Doolittle hydropathy plot of the canonical DAT isoform Q01959. Pg. 18
17. Figure S16. Kyte-Doolittle hydropathy plot of the canonical SERT isoform P31645. Pg. 19
18. Figure S17. Kyte-Doolittle hydropathy plot of the SERT isoform P31645-2. Pg. 20
19. Figure S18. Kyte-Doolittle hydropathy plot of the SERT isoform J3QKP3. Pg. 21

**Table S1. Computationally mapped and predicted potential truncated isoforms of vesicular amine transporter subfamily (VATs).**

| Name | Isoform ID[1] | TM count[2] | Length (aa)[3] | Mass (Da)[4] | Isoelectric point (pI)[5] | GRAVY[6] (cut-off=0) |
|---|---|---|---|---|---|---|
| **NET** | P23975 (Canonical) | 12 | 617 | 69332 | 7.18 | 0.468 |
| | P23975-2 | 12 | 628 | 70548 | 8.44 | 0.414 |
| | P23975-3 | 10 | 512 | 57370 | 6.26 | 0.568 |
| | A0A804HLI4 | 6 | 350 | 39071 | 8.01 | 0.175 |
| | H3BRS0 | 5 | 295 | 33164 | 8.57 | 0.222 |
| **DAT** | Q01959 (Canonical) | 12 | 620 | 68495 | 6.46 | 0.499 |
| **SERT** | P31645 (Canonical) | 12 | 630 | 70325 | 5.89 | 0.422 |
| | P31645-2 | 12 | 672 | 74978 | 5.91 | 0.336 |
| | J3QKP3 | 1 | 72 | 7889 | 5.56 | 1.149 |

[1]The Uniprot entry ID of the isoform (only included isoforms that have length between 10% and 90% of the canonical sequence), P23975-2 and P31645-2 have also been added for comparison purposes. Please see Methods.
[2]Transmembrane (TM) domain count of the protein, derived from the topology information included in the Uniprot entries and Protter.
[3]The amino acid length of the isoform sequence.
[4]Protein mass calculated from sequences via ProtParam.
[4]Protein isoelectric point calculated from sequences via ProtParam.
[5]The GRAVY (Grand Average of Hydropathy) value for the corresponding protein is calculated as the sum of hydropathy values of all the amino acids, divided by the number of residues in the sequence.

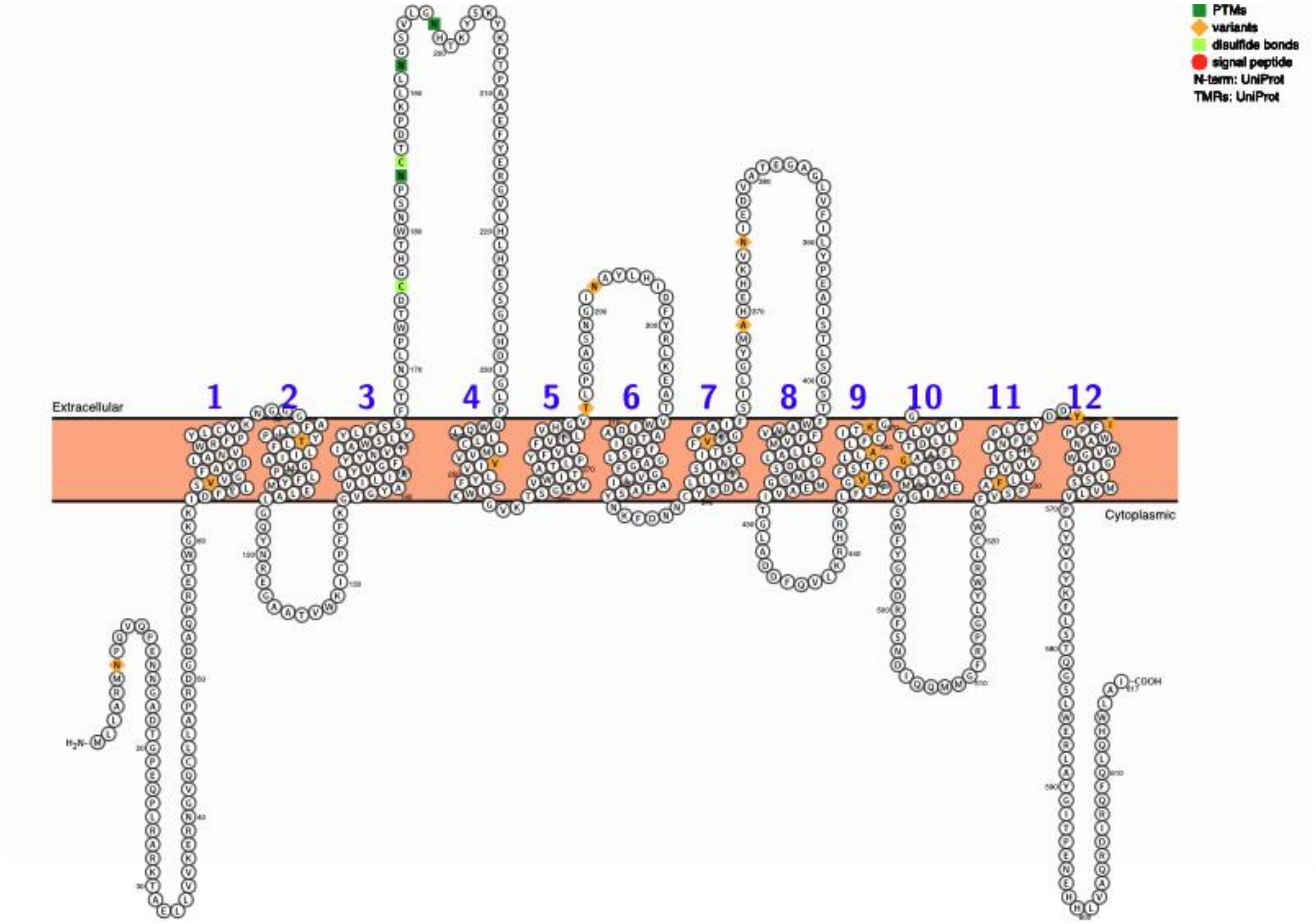

**Figure S1. Sequence and topology visualization of the canonical NET isoform P23975 from Protter.**

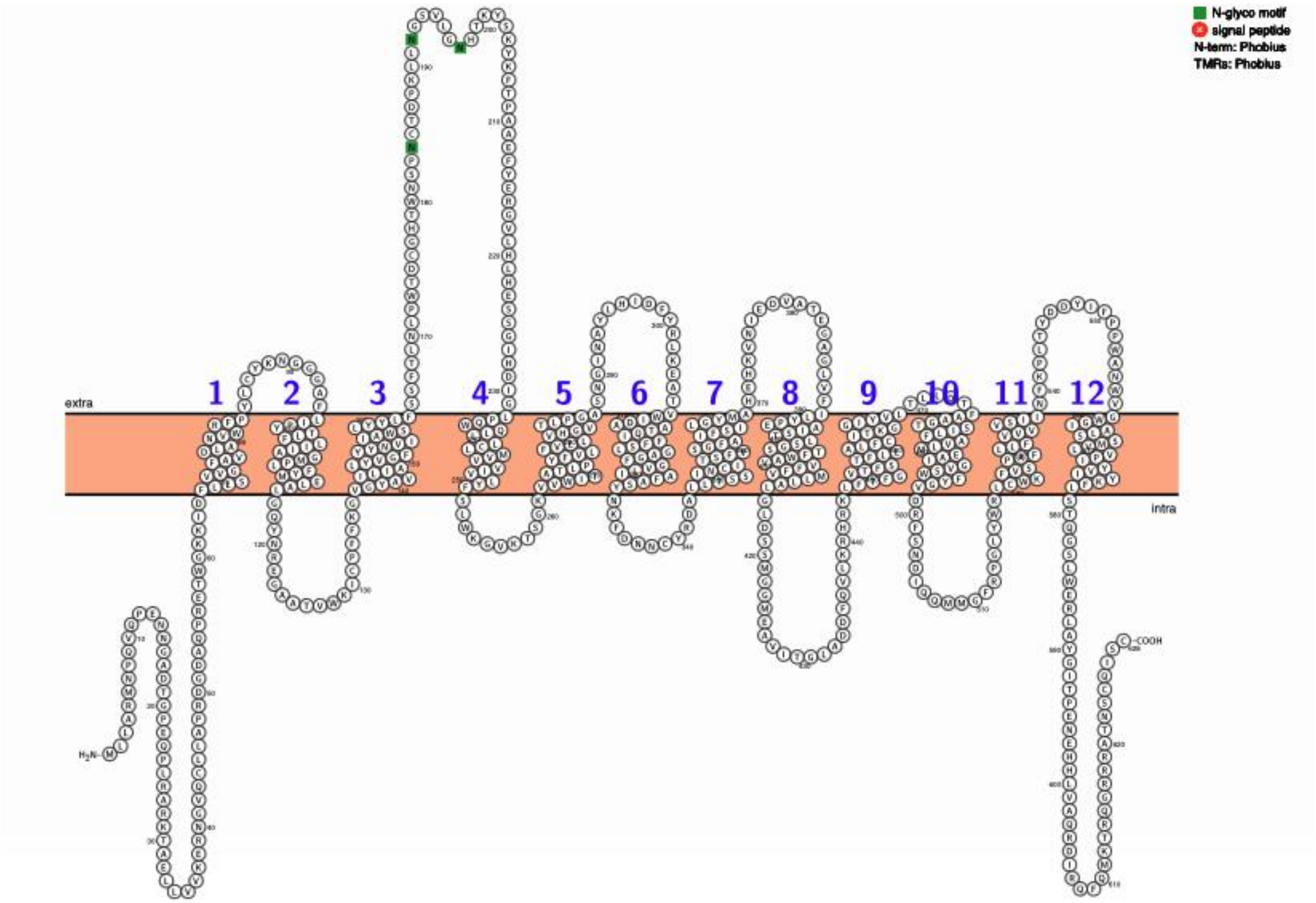

**Figure S2. Sequence and topology visualization of the NET isoform P23975-2 from Protter.**

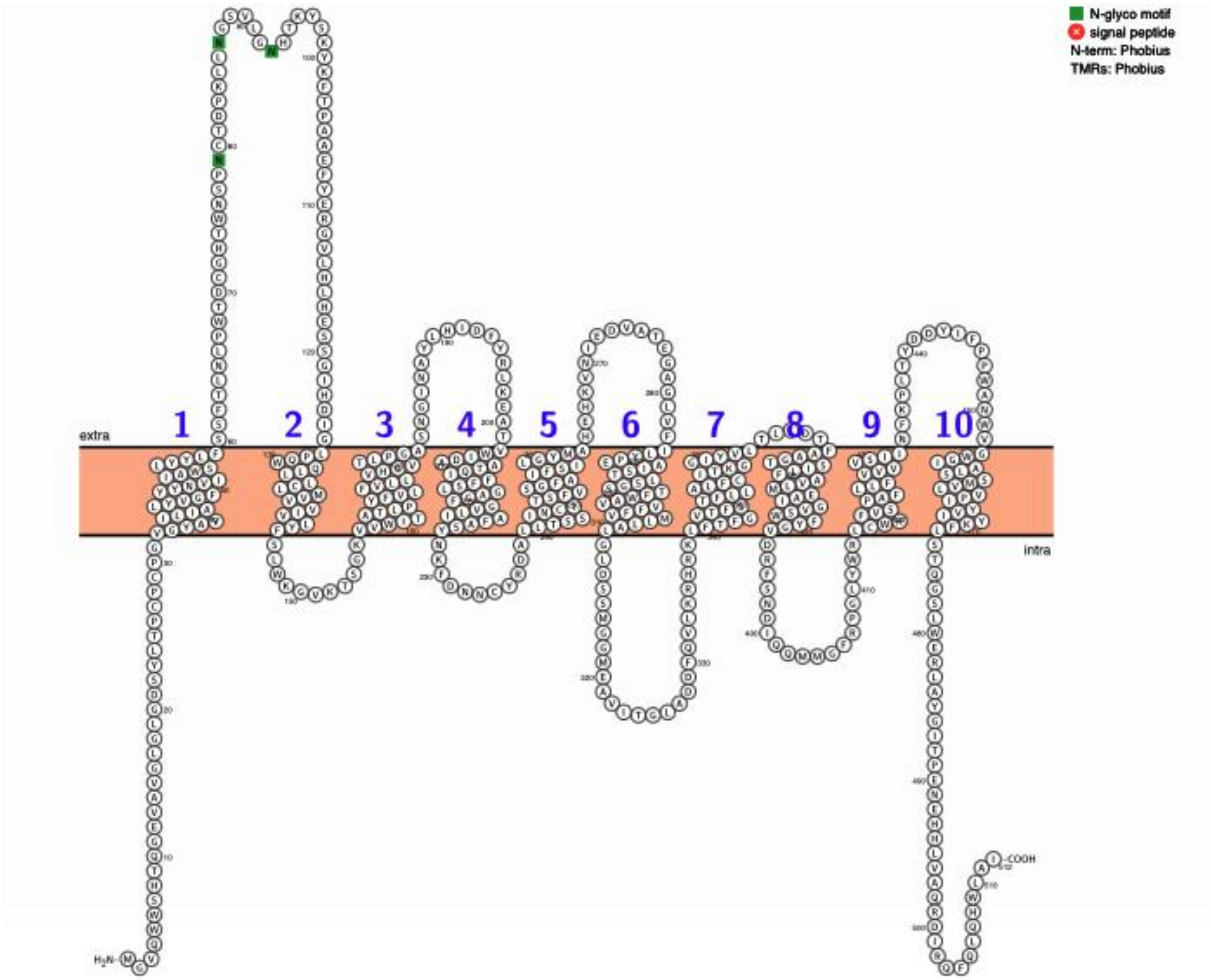

**Figure S3. Sequence and topology visualization of the NET isoform P23975-3 from Protter.**

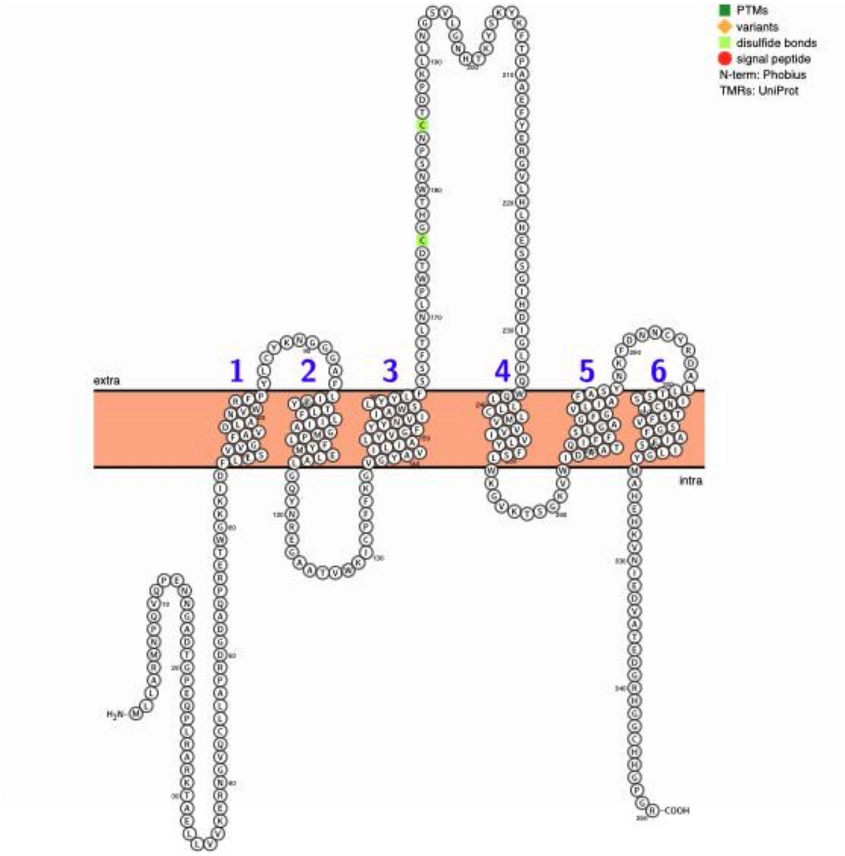

**Figure S4. Sequence and topology visualization of the NET isoform A0A804HLI4 from Protter.**

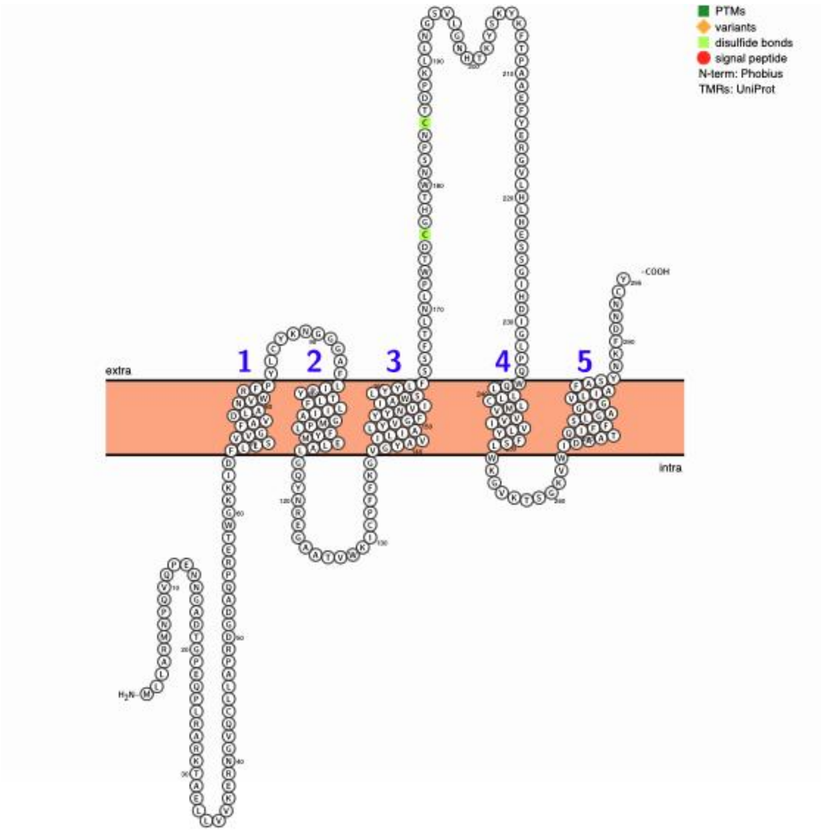

**Figure S5. Sequence and topology visualization of the NET isoform H3BRS0 from Protter.**

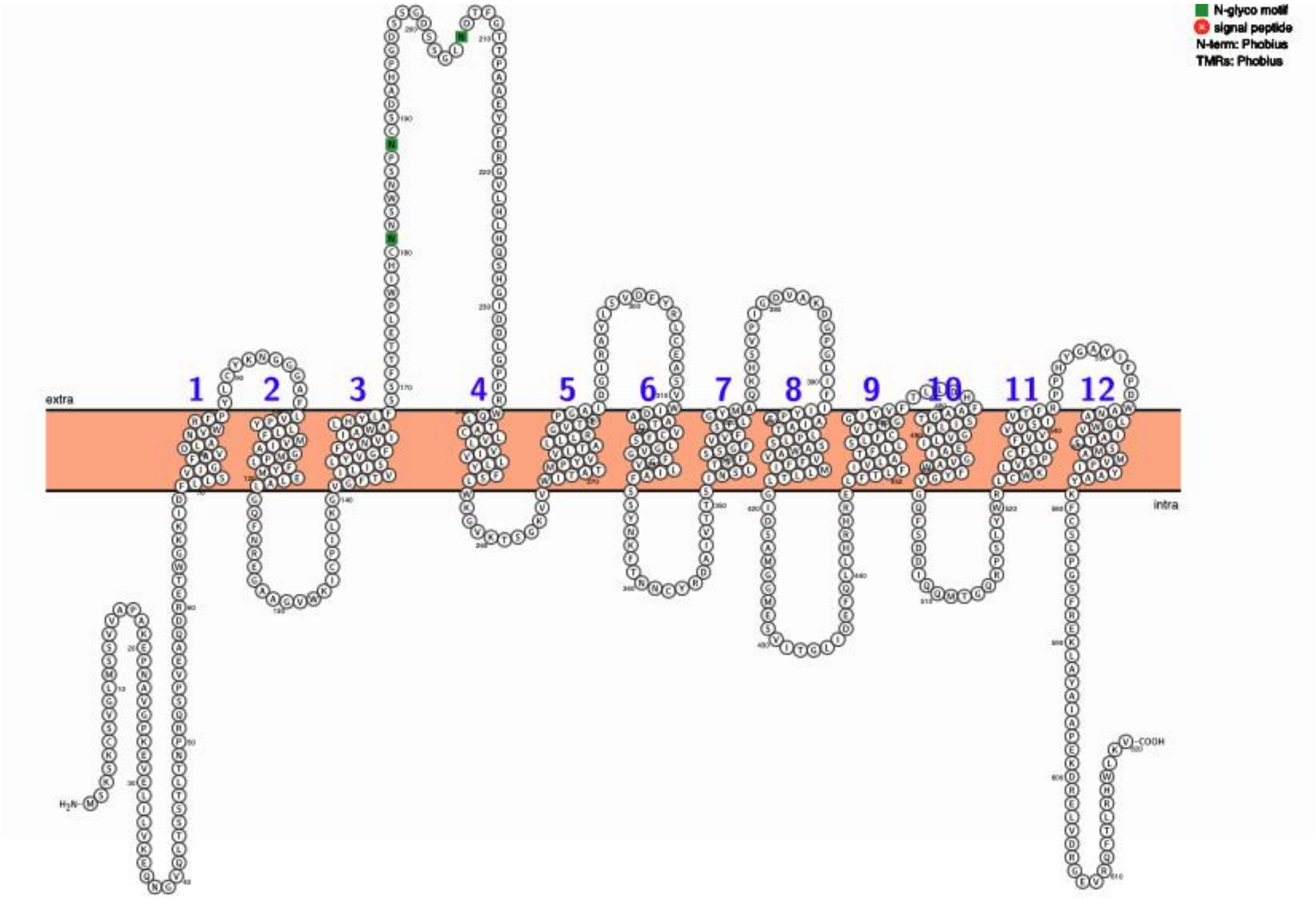

**Figure S6. Sequence and topology visualization of the canonical DAT isoform Q01959 from Protter.**

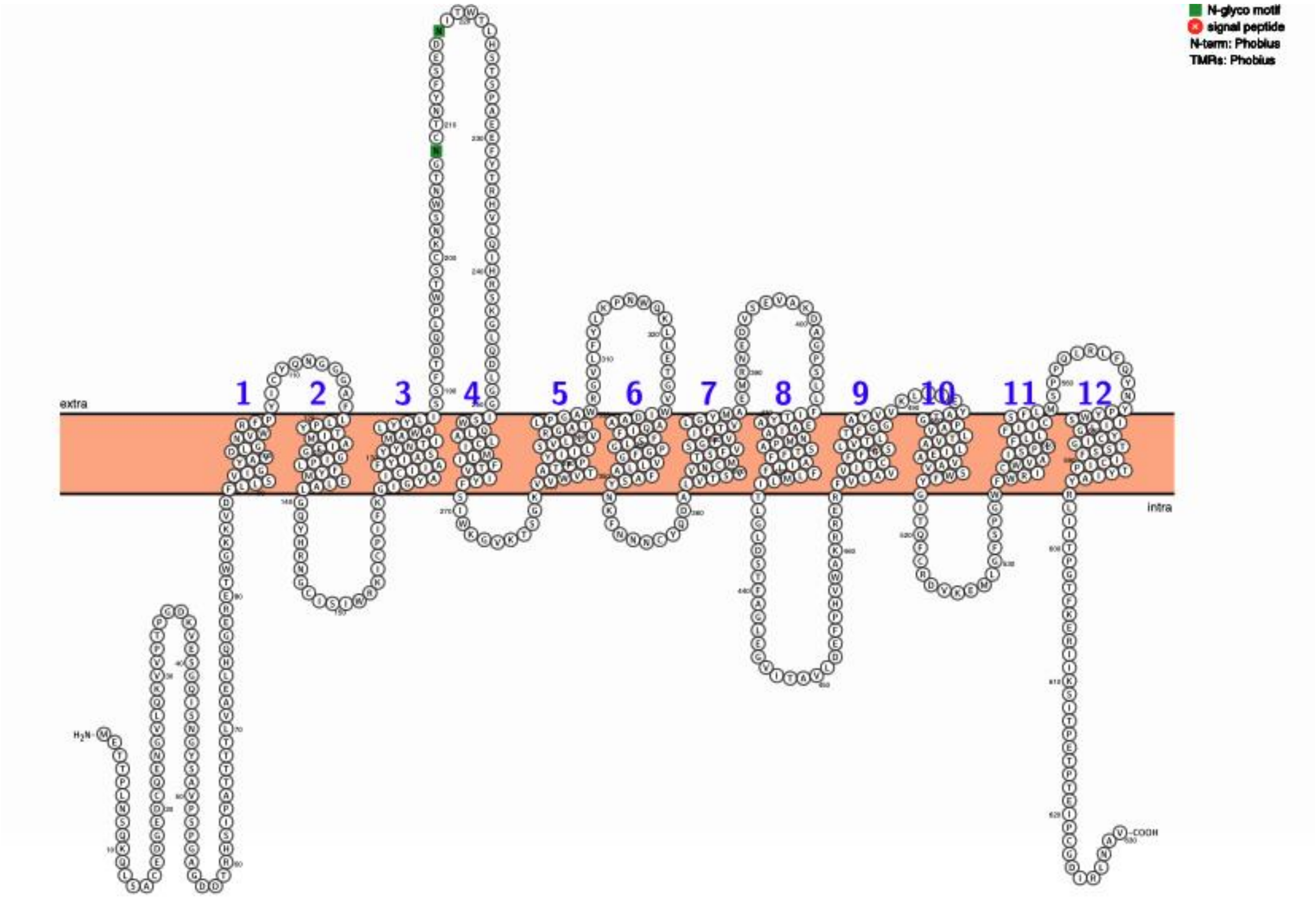

**Figure S7. Sequence and topology visualization of the canonical SERT isoform P31645 from Protter.**

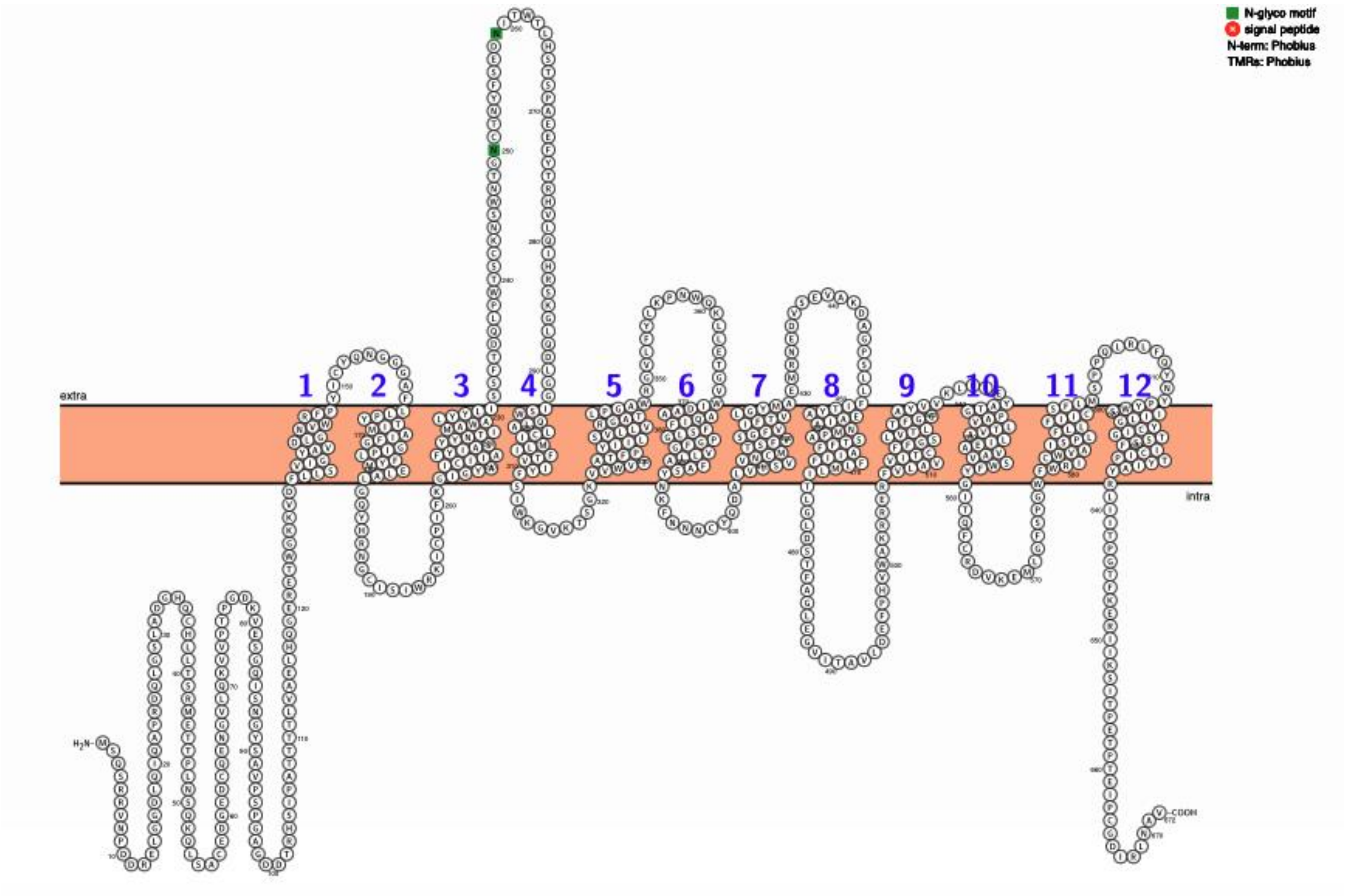

**Figure S8. Sequence and topology visualization of the SERT isoform P31645-2 from Protter.**

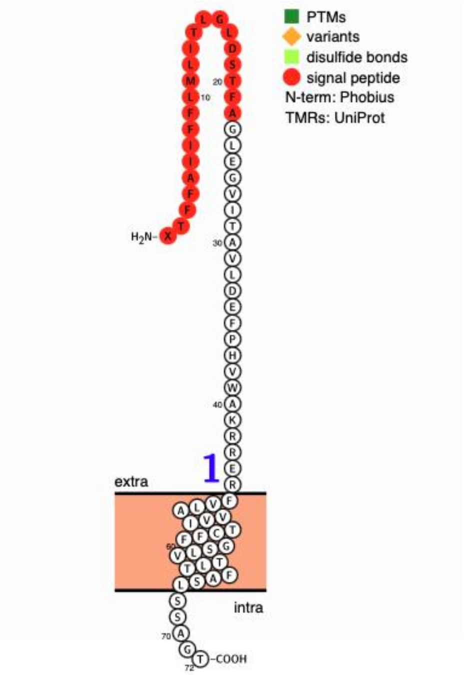

**Figure S9. Sequence and topology visualization of the SERT isoform J3QKP3 from Protter.**

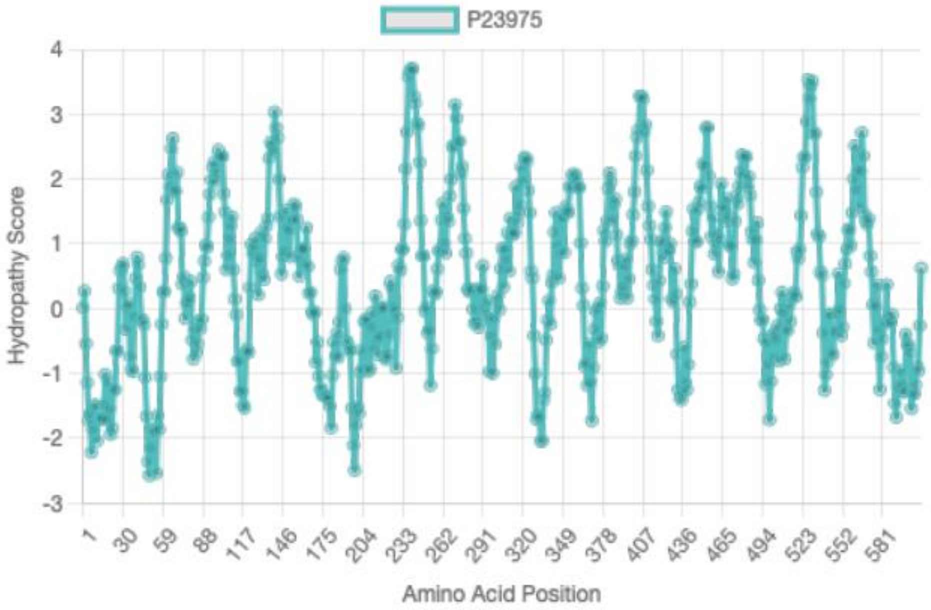

**Figure S10. Kyte-Doolittle hydropathy plot of the canonical NET isoform P23975.**

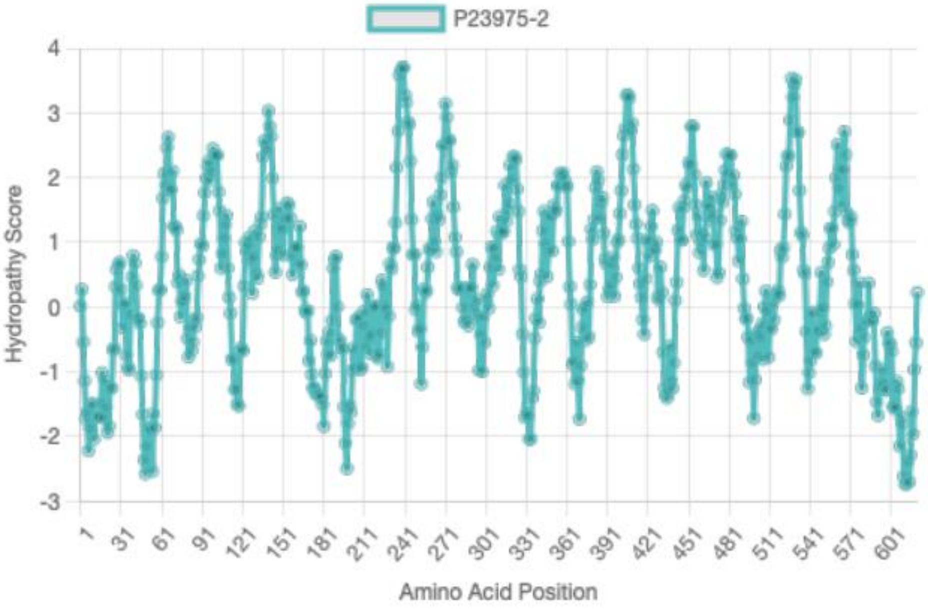

**Figure S11. Kyte-Doolittle hydropathy plot of the NET isoform P23975-2.**

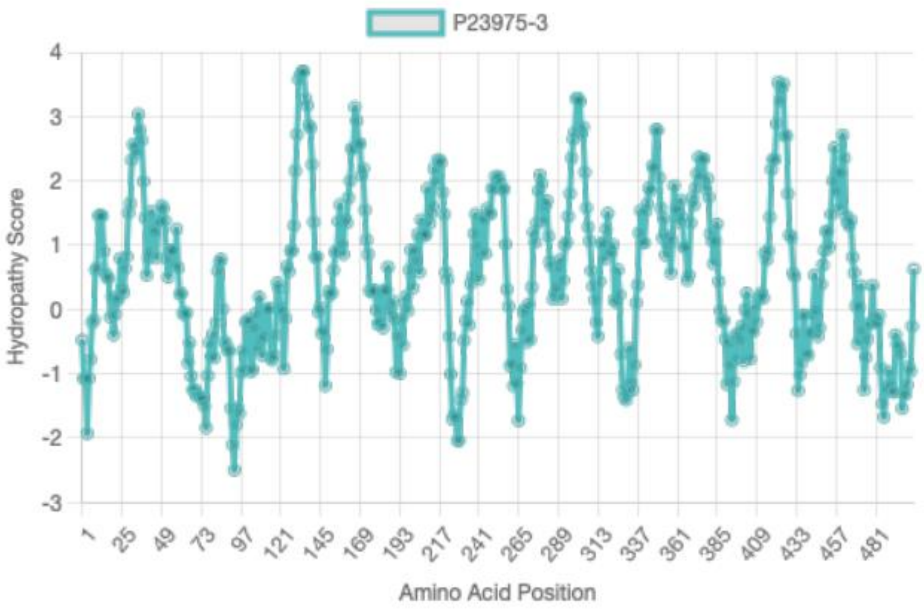

**Figure S12. Kyte-Doolittle hydropathy plot of the NET isoform P23975-3.**

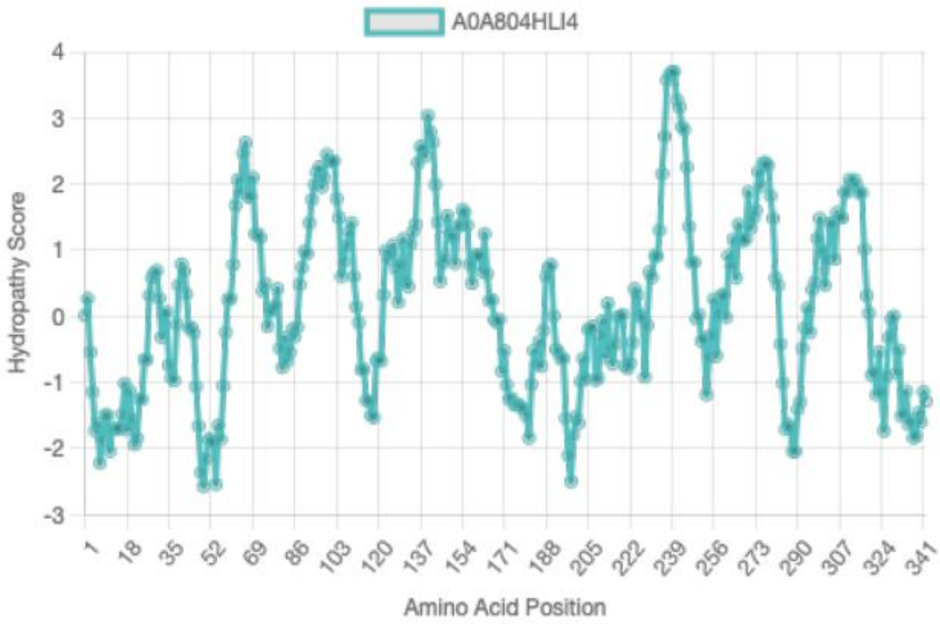

**Figure S13. Kyte-Doolittle hydropathy plot of the NET isoform A0A804HLI4.**

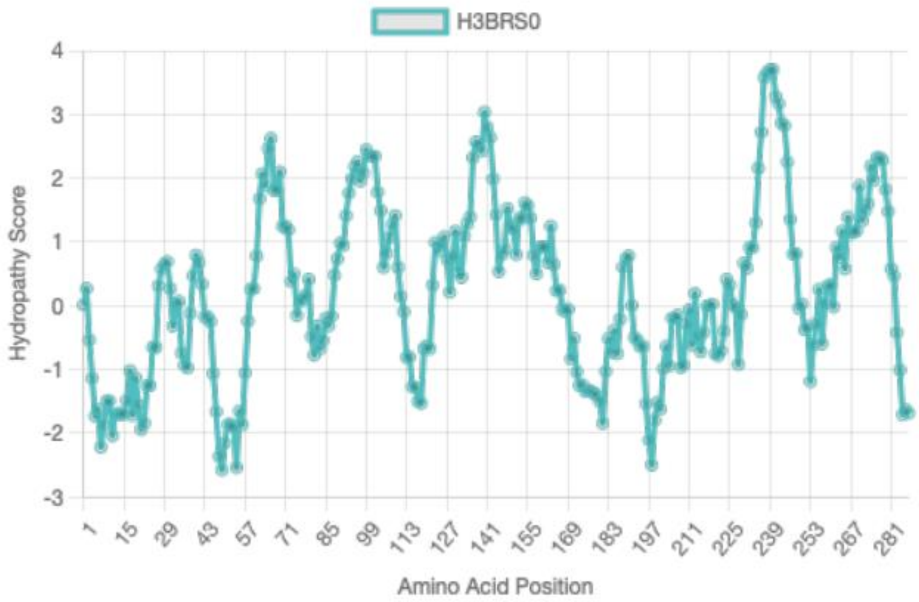

**Figure S14. Kyte-Doolittle hydropathy plot of the NET isoform H3BRS0.**

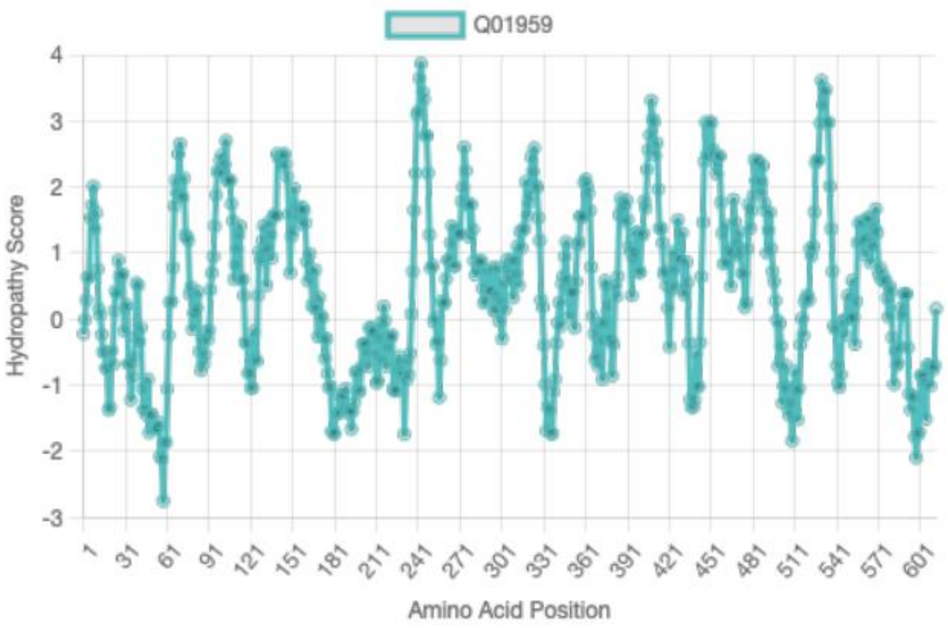

**Figure S15. Kyte-Doolittle hydropathy plot of the canonical DAT isoform Q01959.**

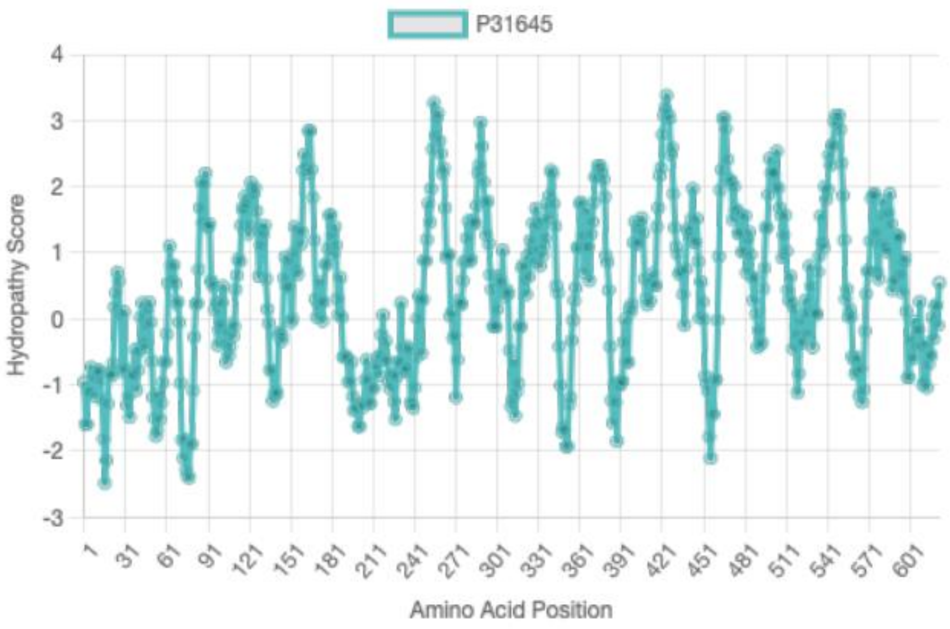

**Figure S16. Kyte-Doolittle hydropathy plot of the canonical SERT isoform P31645.**

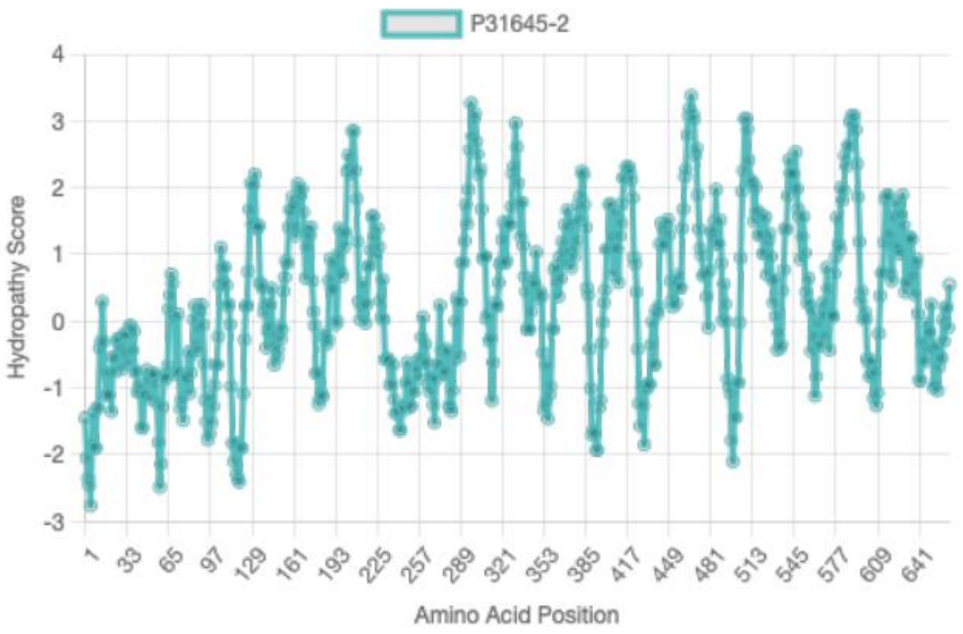

**Figure S17. Kyte-Doolittle hydropathy plot of the SERT isoform P31645-2.**

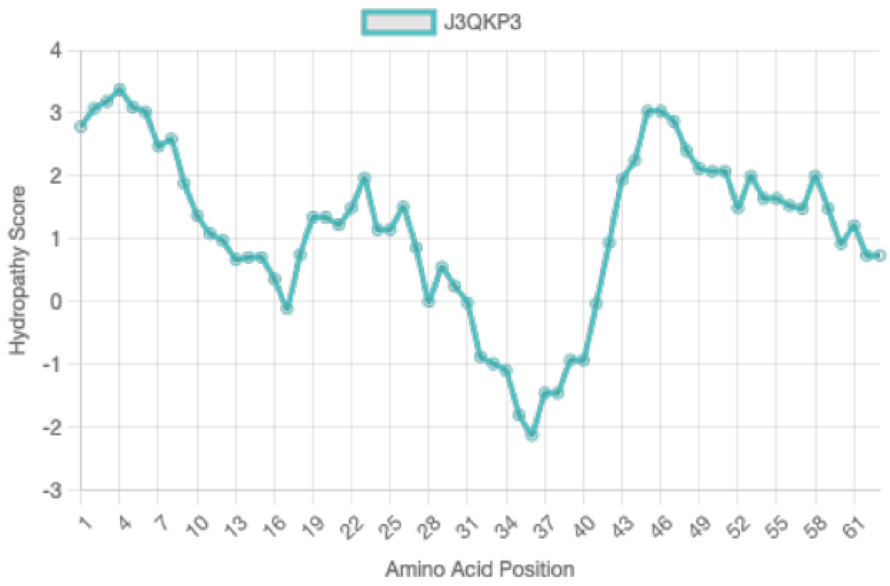

**Figure S18. Kyte-Doolittle hydropathy plot of the SERT isoform J3QKP3.**
